# Curvature Sensing and Membrane Remodeling of the VPS37A N-terminal Domain during Autophagy

**DOI:** 10.1101/2022.11.01.514784

**Authors:** Yansheng Ye, Xinwen Liang, Guifang Wang, Maria C Bewley, Xiaoming Liu, John M. Flanagan, Hong-Gang Wang, Yoshinori Takahashi, Fang Tian

**Affiliations:** Department of Biochemistry and Molecular Biology, The Pennsylvania State University, Hershey, PA USA, 17033; Department of Pediatrics, Division of Pediatric Hematology and Oncology, Pennsylvania State University College of Medicine, Hershey, PA USA, 17033

## Abstract

VPS37A, a component of ESCRT-I, is essential for recruiting a subset of ESCRT proteins that seal the phagophore during autophagosome biogenesis. In this study, we uncover two hydrophobic motifs in the VPS37A N-terminal 148 amino acids (VPS37A^1-148^) that selectively interact with highly curved membranes. Mutations in these motifs nearly abolish VPS37A membrane binding *in vitro* and compromise its localization to the phagophore and autophagic flux *in vivo*. We also determined the solution structure of residues 21 to 131 and demonstrated that it is the UEVL (ubiquitin E2 variant-like) domain. Intriguingly, this domain remodels highly curved liposomes to high-order structures. We suggest that the specific interactions between VPS37A^1-148^ and the curved membrane may facilitate the recruitment of VPS37A to the phagophore and its subsequent closure. Our results support the premise that the distinct membrane architecture of the cup-like phagophore spatiotemporally regulates autophagosome biogenesis.

## Introduction

During macroautophagy (hereafter referred to as autophagy), cytoplasmic components targeted for degradation are sequestered into a double-membrane vesicle (autophagosome) prior to lysosomal degradation. The biogenesis of a functional autophagosomes occurs via a specific and extensive membrane remodeling sequence that includes initiation, nucleation of the isolation membrane, phagophore expansion, and phagophore closure^1–3^. While many of these steps are coordinated by Atg (autophagy-related) proteins^4^, we recently discovered that several components of the endosomal sorting complexes required for transport (ESCRT) machinery, including the human proteins VPS37A, VPS28, TSG101, CHMP2A, and VPS4, are necessary for sealing the cup-shaped phagophore and forming the resulting autophagosome^5–7^. Specifically, VPS37A recognizes the phagophore and recruits a subset of ESCRT components to complete phagophore closure. Importantly, deletion of the first 90 residues of VPS37A abolishes its localization to the phagophore but has little effect on its role in the degradation of the epidermal growth factor receptor in the multivesicular body (MVB) pathway^5^. These results suggest that the VPS37A N-terminal serves a specific function during phagophore closure but the exact nature of this process is unclear.

The ESCRT pathway is highly conserved and is required for the formation of multivesicular bodies, cytokinesis, plasma membrane and lysosome repair, nuclear pore reformation, exosome biogenesis, autophagy, and virus budding^8–10^. The ESCRT machinery draws opposing membranes together and mediates the final membrane scission reaction required for processes such as vesicle inward budding into an endosome and virus budding from the plasma membrane. These features are common mechanistic elements of phagophore closure. VPS37A is a subunit of the heterotetrameric ESCRT-I complex. In the crystal structure of the yeast ESCRT-I complex, its core consists of a flatted headpiece linked to a long stalk formed by Vps23, Vps28, Vps37, and Mvb12^11^. The N-terminal ubiquitin E2 variant (UEV) domain of Vps23 (TSG101 in human) and the C-terminal domain of Vps28 are flexibly tethered to the core for targeting ubiquitinated cargos and recruiting ESCRT-II, respectively. The core structure of human ESCRT-I formed by TSG101-VSP28-VPS37B-MVB12A was recently determined to be structurally homologous to yeast’s^7^.

Human VPS37A is a multidomain protein containing 397 residues. The C-terminal sequence contains a conserved Mod(r) domain which is part of the core structure of the ESCRT-I complex (Extended Data Fig. 1a). The first ~220 residues of VPS37A are unique among its four mammalian homologs (VPS37A-D, Extended Data Fig. 2). This region is predicted to consist of a ubiquitin E2 variant-like (UEVL) domain (previously referred to as a putative ubiquitin E2 variant domain^5^), and long stretches of unstructured regions that are poorly characterized (Extended Data Fig. 1a). Here we identify two hydrophobic motifs in the unstructured regions that selectively interact with highly curved membranes *in vitro*, and characterize them functionally. Mutations in these two regions result in defective VPS37A phagophore localization and autophagy flux *in vivo*. In addition, NMR structural studies indicate that residues 23 to 131 form a UEVL domain. Surprisingly, unlike other UEV domaincontaining proteins^12, 13^, the VPS37A UEVL domain does not interact with monoubiquitin (Ub). Instead, it remodels highly curved liposomes to high-order structures. While factors that recruit VPS37A to the phagophore are unknown, we speculate that the preference of its N-terminus for interacting with curved membranes may lead or contribute to VPS37A localization to the highly curved membrane edge of the phagophore, and facilitate the dynamic membrane remodeling of the phagophore and autophagosome biogenesis during autophagy.

## Results

### VPS37A N-terminal performs unique function(s) during autophagy

Our previous study has shown that the N-terminal of VPS37A is required for its phagophore localization^5^. However, since the putative UEVL domain in the VPS37A N-terminus does not have notable sequence homology with any proteins of known structure, we used NMR to determine its structure. Based on a previous biological study^14^, we initially used a construct consisting of residues 1 to 148 (VPS37A^1-148^) that is predicted to contain the putative UEVL domain (Extended Data Fig. 1a and 1b). Using ^13^C and ^15^N labeled proteins and triple resonance experiments, we assigned all of the resonances from non-proline residues except S24, H89, and N-terminal 13 residues (BMRB: 51558). Since there are only five unassigned peaks in the 2D TROSY spectrum (Extended Data Fig. 1c), resonances from the majority of the first 13 residues were not observed, likely due to exchange broadenings. In addition, analyses of secondary chemical shifts of ^13^C_α_ and ^13^C_β_ for residues 14 to 20 indicate that this region is unstructured (data not shown). Therefore, to facilitate NMR structure determination we prepared a construct comprising residues 21 to 148 of VPS37A (VPS37A^21-148^, Extended Data Fig. 1b); its 2D ^15^N-^1^H correlation spectrum superimposes with that of the initial VPS37A^1-148^ construct in Extended Data Fig. 3. Minor shifts in these two spectra indicate that removing the N-terminal first 20 residues does not perturb the structure. The solution structure of VPS37A^21-148^ determined with NOEs, RDCs, and torsion angle restraints is shown in Fig. 1a (PDB ID: 8E22, BMRB ID:31039, Table S1). As expected, residues 21 to 131 adopt the characteristic α/β fold found in canonical UEV domains, while residues 132 to 148 are unstructured.

**Fig. 1:**
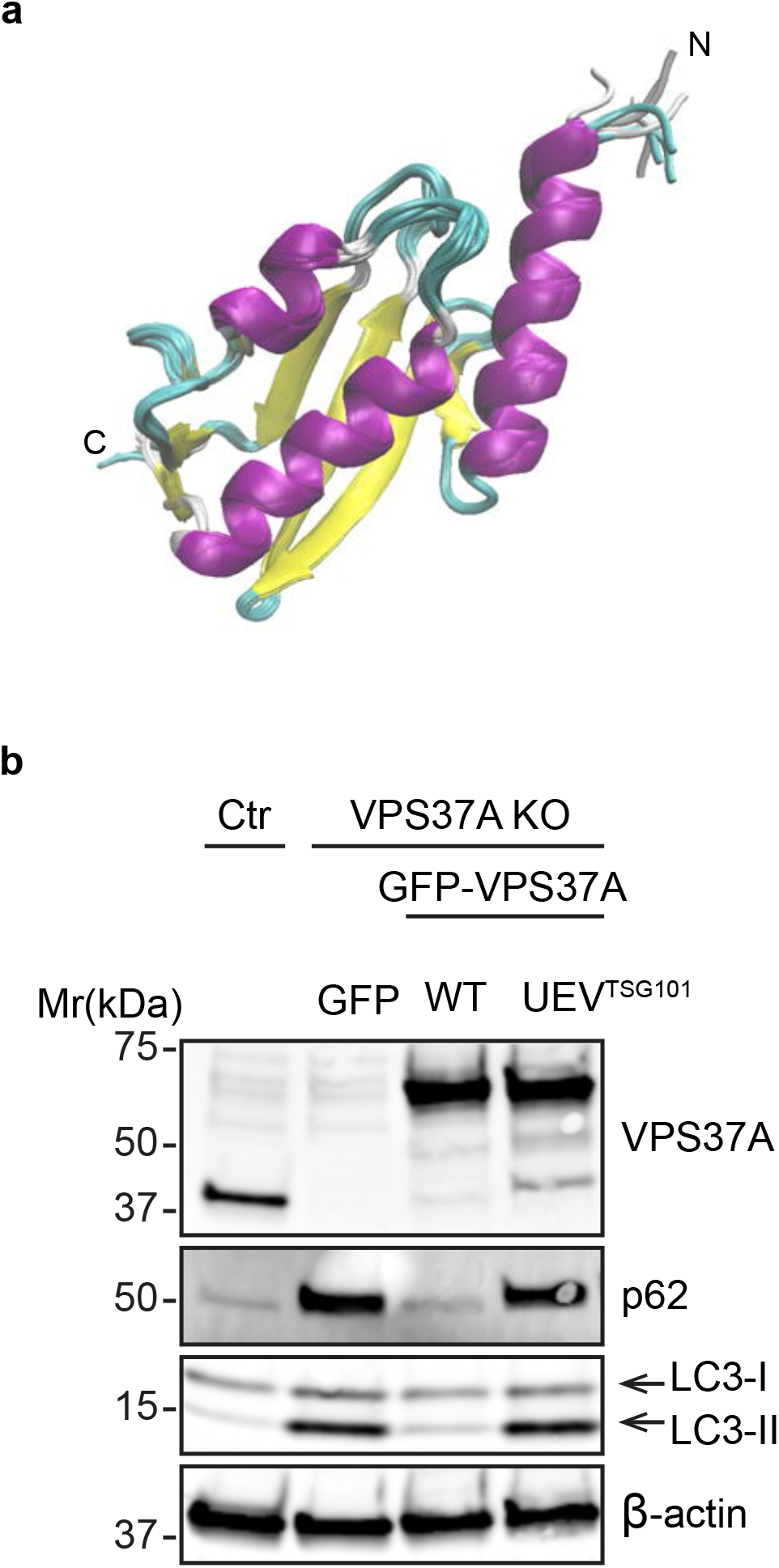
Residues 21 to 148 of VPS37A form a UEVL domain that cannot be functionally substituted by TSG101 UEV. a. Overlay of ten lowest-energy NMR structures of VPS37A^21-148^. Unstructured region of residues 133 to 148 is not shown. N and C indicate N- and C- terminals, respectively. b. Immunoblot analysis of VPS37A KO U-2 OS cells expressing the indicated constructs (GFP-only, GFP-VPS37A wild-type (WT), and GFP-UEV^TSG101^-VPS37A (UEV ^TSG101^)) relative to the control (Ctr).

As the core structure of the VPS37A UEVL domain is similar to those of other UEV proteins (Extended Data Fig. 4a), we tested whether UEV domain of TSG101 can functionally replace that of VPS37A. Using a structure-based sequence alignment as a guide (Extended Data Fig. 4b), we constructed a N-terminal GFP-tagged protein in which residues 1 to 149 of VPS37A were replaced with the corresponding sequence (residues 1 to 145) of TSG101 (GFP-UEV^TSG101^-VPS37A, Extended Data Fig. 1d). Transduction of VSP37A KO cells with wild-type GFP-VPS37A rescued the autophagic defect of the deletion cell lines and restored the levels of the autophagosomal membrane marker LC3-II and the autophagic substrate p62 to those of control wild-type cells (Fig. 1b). By contrast, transduction of GFP-UEV^TSG101^-VPS37A into VPS37A KO cells failed to rescue the autophagic defects as judged by equivalent levels of LC3-II and p62 accumulation between GFP-only and GFP- UEV^TSG101^-VPS37A transductions (Fig. 1b). The fact that this N-terminal region of VPS37A is not functionally interchangeable with the TSG101 UEV *in vivo* points to its novel but uncharacterized role(s) in phagophore sealings.

### VPS37A N-terminal selectively interacts with highly curved membranes

Yeast Vps37 (yVps37) has an N-terminal basic helix (residues 1-21, H_0_) which interacts with acidic lipids. Since deletion of H_0_ abolishes the binding of yeast ESCRT-I to the membrane, this helix has been suggested as the driving force for the recruitment of yeast ESCRT-I to the membrane^11^. This observation led us to investigate whether human VPS37A^1-148^ interacts with the membrane. In liposome flotation assays most of the VPS37A^1-148^ protein was detected in the top (T) lipid-containing fraction with sonicated liposomes consisting of POPC:DOPG:DOPE in a molar ratio of 3:2:5 (20.0 nm average radius measured by dynamic light scattering, DLS) as shown in Fig. 2a and 2b. However, with three different preparations of larger liposomes (average radii of 38.5, 48.9 and 69.3 nm determined by DLS and prepared by extrusion with 50, 100, and 200 nm filters, respectively), the protein was predominantly detected in the aqueous solution fraction (bottom, B), indicating that the protein minimally binds to larger and less curved liposomes. Thus, VPS37A^1-148^ interacts with model membranes, and its interaction depends on vesicle size; VPS37A^1-148^ clearly shows a preference for small lipid vesicles with highly curved membranes. By contrast, the TSG101^1-145^ does not float with or bind to any of these liposome preparations (Extended Data Fig. 5).

**Fig. 2:**
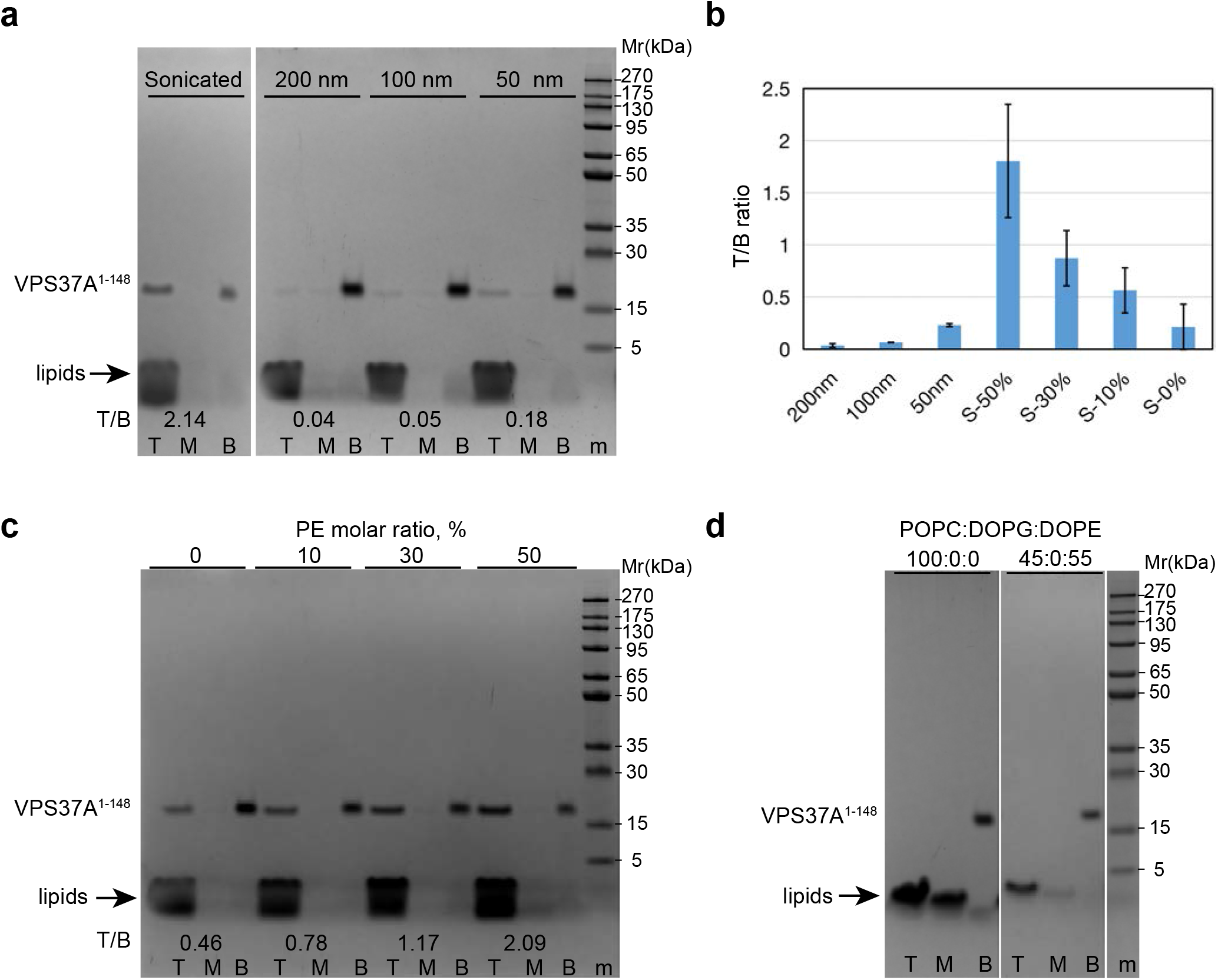
VPS37A^1-148^ selectively interacts with highly curved membranes. a. Gel images of liposome flotation assays for VPS37A^1-148^ mixed with sonicated or extruded liposomes with membrane pore sizes of 50, 100 or 200 nm. (POPC:DOPG:DOPE=3:2:5, protein:lipid=1:400, molar ratio). T, M and B represent top, middle and bottom layers after centrifugation. Protein marker is indicated by m. Amount of VPS37A^1-148^ in the top layer to that in the bottom (T/B) is quantitated by ImageJ. b. Plots of VPS37A^1-148^ in top layer versus bottom layer for extruded liposomes (POPC:DOPG:DOPE=3:2:5) with membrane pore sizes of 50, 100, or 200 nm, and sonicated liposomes containing 0, 10, 30, or 50% PE (molar ratio, referred to as S-0%, S-10%, S-30%, and S-50%, respectively). Data are presented as mean ± SD (standard deviation). Quantifications obtained from three separate measurements (n= 3). c. Gel images of liposome flotation assays for VPS37A^1-148^ in sonicated liposomes consisting of 20% DOPG and 0, 10, 30, or 50% of PE (molar ratio). Their plots are shown in b. d. Gel images of liposome flotation assays for VPS37A^1-148^ in sonicated liposomes containing 0 or 55% of DOPE (molar ratio) without negatively charged PG.

Hydrophobic insertion is one of the major molecular mechanisms responsible for membrane curvature recognition^15, 16^. Packing defects in the outer leaflet of the lipid bilayer are a hallmark of highly curved membranes, and curvature-sensitive molecules preferentially embed into surfaces that display this feature. By extension, introducing packing defects into the bilayer by incorporating lipids with small headgroups, such as PEs, should strengthen the membrane binding of curvature-sensitive molecules^17^. To determine how VPS37A^1-148^ targets small, highly curved membranes, we repeated the liposome flotation assay with sonicated liposomes containing different percentages of PE lipids. As shown in Fig. 2b and 2c, we found that the amount of VPS37A^1-148^ in the top fraction correlates with increasing percentages of PE lipids in liposomes. This suggests that the membrane binding of VPS37^1-148^ is enhanced by PE lipids, consistent with bilayer packing defects contributing to the enhanced binding. In addition, VPS37^1-148^ does not float with or bind to liposomes without DOPG lipids (Fig. 2d), indicating that its membrane binding requires negatively charged lipids. Together, these results are consistent with a model in which VPS37^1-148^ selects curved membranes by hydrophobic insertion and both hydrophobic and electrostatic interactions stabilize binding to the membrane.

To identify the sequence(s) responsible for the membrane curvature-sensitive interaction of VPS37^1-148^, we used bicelles as a membrane model and characterized the interactions by high-resolution NMR. We have recently shown that bicelles are an effective model for highly curved membranes in the study of human Atg3 because the dynamic planar surfaces of bicelles are loosely packed and presumably mimic the type of packing defects found in highly curved membranes^18^. An overlay of 2D ^15^N-^1^H correlation spectra of VPS37A^1-148^ in the presence and absence of bicelles is shown in Fig. 3a. These spectra were acquired at 15 °C to increase the lifetime of VPS37^1-148^ NMR resonances in bicelles, which generally disappear within an hour at room temperature (data not shown). In the presence of the bicelles several peaks were shifted including resonances from residues A136 to N143, the indole amide proton of residue Trp3, and five unassigned resonances (most of which are presumably from 13 unassigned residues of the N-terminal disordered region as described above). Based on sequence alignment (Extended Data Fig. 6), these perturbed residues are cluster around two highly conserved regions containing bulky hydrophobic amino acids, ^3^WLFP and ^137^FPYL (highlighted in yellow), which straddle the UEVL domain. These two regions are of particular interest since insertion of a hydrophobic moiety into one leaflet of lipid bilayers is a common mechanism for membrane curvature recognition and generation. To examine the relative contributions of each region to membrane binding, we replaced the sequence F^137^PYL with alanine residues in the VPS37A^1-148^ and VPS37A^21-148^ (without the ^3^WLFP motif) constructs referred to hereafter as VPS37A^1-148, 4A^ and VPS37A^21-148, 4A^ (Extended Data Fig. 1b). Fig. 3b and 3c show the results of liposome flotation experiments for these variants. Compared with VPS37A^1-148^, the ratios of membrane-bound to free of VPS37A^21-148^ and VPS37A^21-148, 4A^ were reduced by ~60% and the ratio of membrane-bound to free of VPS37A^1-148, 4A^ was reduced by ~25%. These observations suggest that both hydrophobic motifs contribute to membrane interactions, although the loss of the ^3^WLFP motif has a more significant effect.

**Fig. 3:**
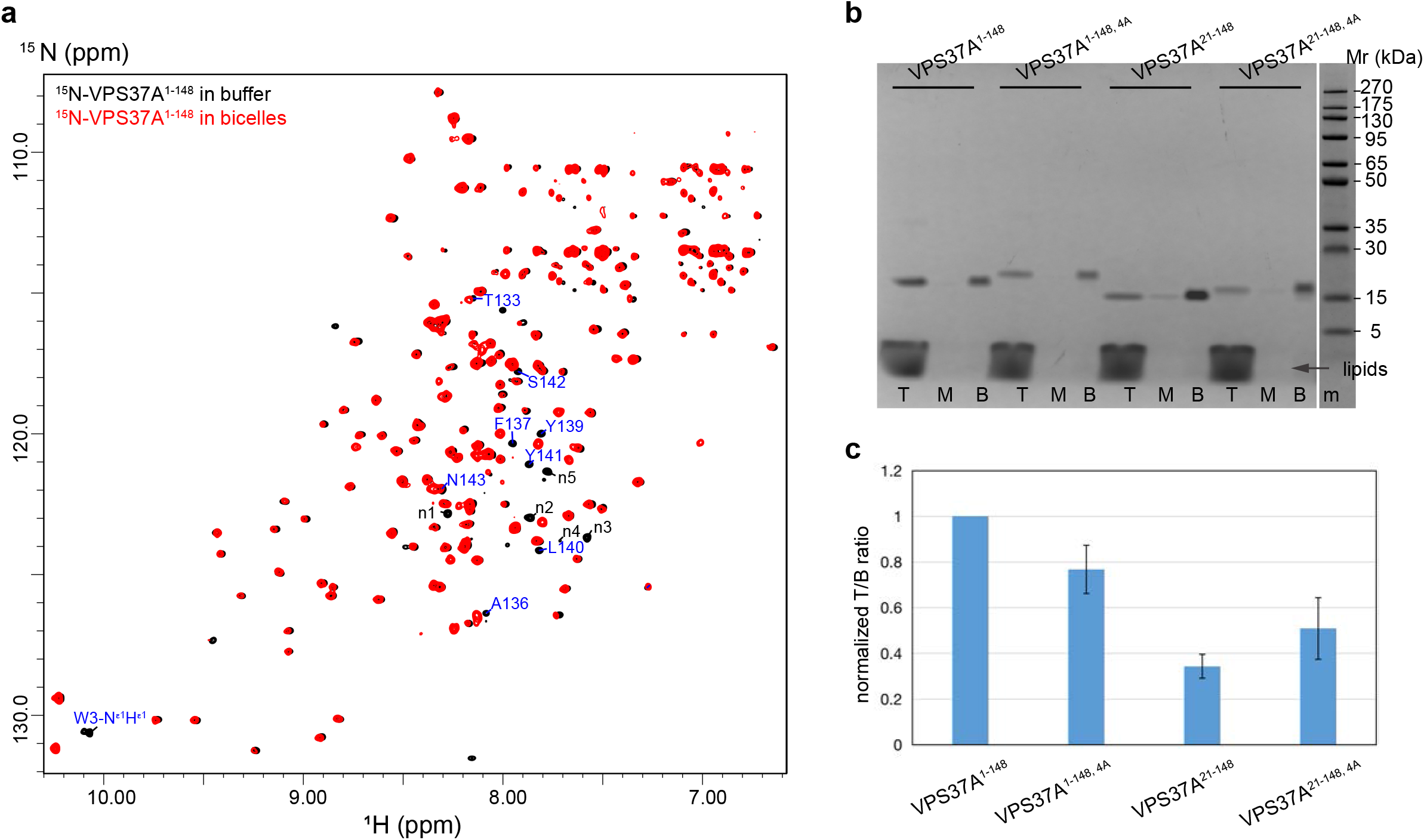
Two conserved hydrophobic motifs in VPS37A^1-148^ interact with the membrane. a. Overlay of ^15^N-labeled VPS37A^1-148^ TROSY spectra in the absence (black) and presence (red) of bicelles (DMPC:DMPG:DHPC = 4:1:10, molar ratio, q=0.5). Several perturbed resonances are labeled with their assignments. Unassigned resonances are labeled with n1 to n5. b. Gel images of liposome flotation assays for VPS37A^1-148^, VPS37A^1-148, 4A^, VPS37A^21-148^, and VPS37A^21-148, 4A^ mixed with sonicated liposomes (POPC:DOPG:DOPE=3:2:5, protein:lipid=1:400, molar ratio). T, M and B represent top, middle and bottom layers after centrifugation. Protein marker is indicated by m. c. Plots of VPS37A^1-148^ and its mutants in the top relative to bottom layer in flotation experiments using sonicated liposomes (POPC:DOPG:DOPE=3:2:5). The T/B ratio of mutants is normalized to the ratio of VPS37A^1-148^. Amount of protein in top layer relative to bottom (T/B) is quantitated by ImageJ. Data are presented as mean ± SD from three independent experiments (n=3 for each construct).

### Membrane curvature-sensitive binding facilitates targeting of VPS37A to phagophores for autophagosome completion

Having established that the two hydrophobic sequences are involved in membrane binding *in vitro*, we evaluated their effect in ESCRT-I targeting during autophagy by replacing endogenous VPS37A with exogenously expressed GFP-tagged wild-type VPS37A or a VPS37A mutant of which the first 20 residues were removed and with F^137^PYL replaced by A^137^AAA (Mut, VPS37A^Δ20, 4A^). To prevent VPS4-dependent ESCRT disassembly, the cells were transfected with siRNAs targeting the ESCRT-III component CHMP2A. The successful depletion of CHMP2A was verified by immunoblotting (Extended Data Fig. 7). Consistent with our previous findings^5, 7^, siCHMP2A-transfected cells accumulated GFP-VPS37A and pHuji-LC3-double-positive (the phagophore/nascent autophagosome marker) structures^19^. This indicates that, in the absence of CHMP2A, ESCRT-I binding to phagophores is stabilized when VPS4 recruitment was inhibited. The accumulation of GFP-VPS37A and pHuji-LC3 foci were also observed in the mutant-expressing cells upon CHMP2A depletion (Fig. 4a and 4b). However, the level of VPS37A^Δ20, 4A^ accumulation was much lower than wild-type VPS37A. Moreover, pHuji-LC3-labelled structures in the mutant-expressing cells were frequently negative for VPS37A signals, suggesting an attenuation of ESCRT-I membrane targeting during autophagy.

**Fig. 4:**
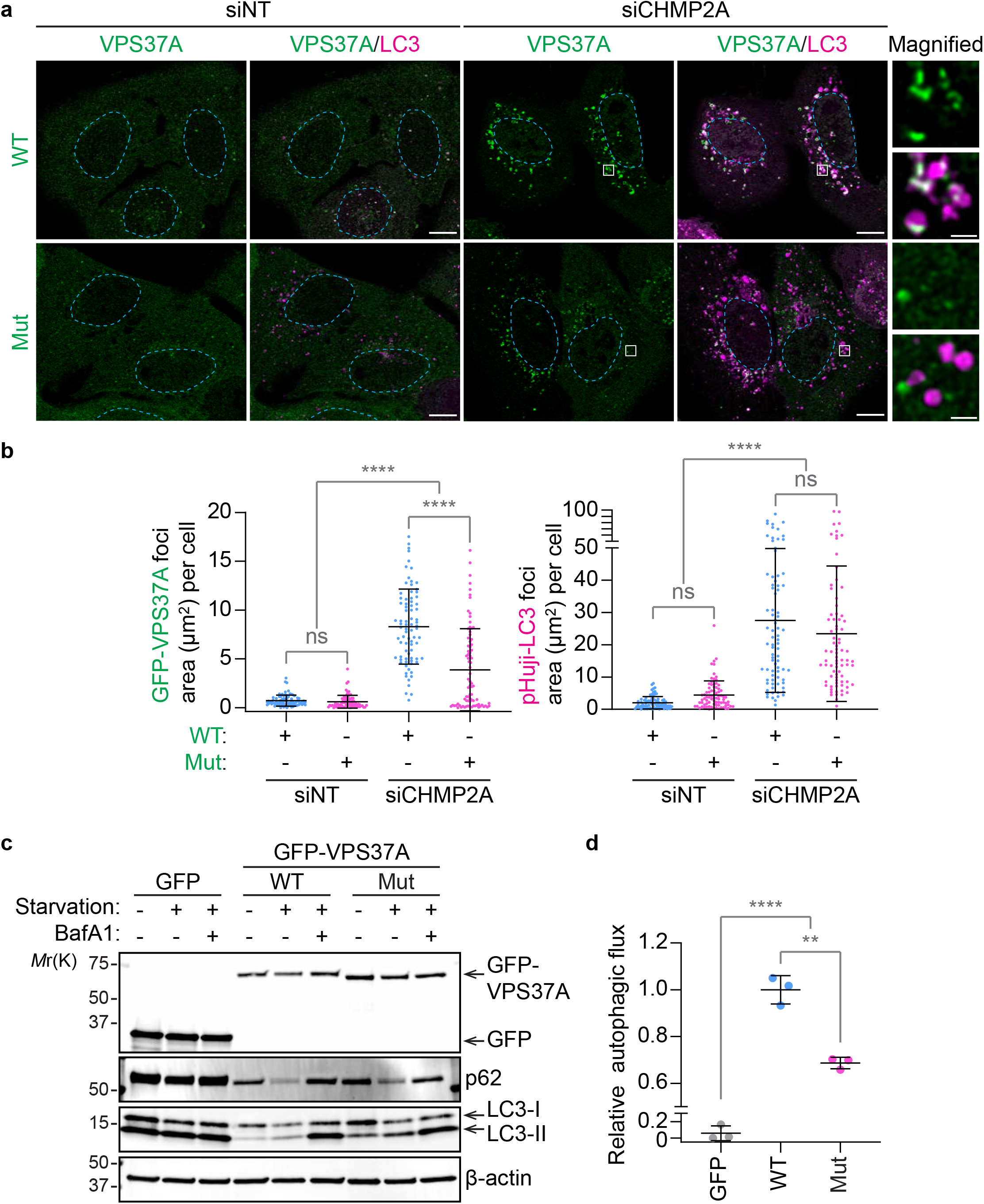
VPS37A^1-148^ membrane binding promotes ESCRT-I targeting to autophagosomal membranes. a. Confocal images of VPS37A KO U-2 OS cells that were stably transduced with wild-type VPS37A (WT) or its mutant without the first N-terminal 20 residues and with F^137^PYL to A^137^AAA mutation of VPS37A (Mut, VPS37A^Δ20, 4A^), transfected with CHMP2A siRNA (siCHMP2A) or control non-targeting siRNA (siNT) for 40 hrs, and starved for 2 hrs. Representative images from 2 independent experiments are shown. The scale bars represent 10 μm, and 1 μm in the magnified images. b. Fluorescence intensities of GFP-VPS37A and the phagophore/nascent autophagosome marker pHuji-LC3-positive foci per cell area in A were quantified and shown (n = 80 cells). ns, not significant; ****, p ≤ 0.0001. c. Immunoblot analysis of the indicated cells that were starved in the presence or absence of 100 nM BafA1 for 3 hrs. d. Autophagic flux under starvation conditions calculated as described in the Methods section was normalized to the mean of the control WT cells (n = 3 independent experiments).

We next examined the functional significance of VPS37A lipid binding in autophagy. In contrast to the expression of GFP-tagged wild-type VPS37A, expression of the GFP-tagged VPS37A^Δ20, 4A^ mutant in VPS37A KO cells resulted in the accumulation of both the autophagosomal membrane marker LC3-II and the autophagic substrate p62, regardless of nutrient status and independent of the lysosomal inhibitor Bafilomycin A1 (BafA1). This indicates an impairment of autophagic flux (Fig. 4c and 4d). Collectively, these results suggest that membrane curvature sensing, mediated by the two hydrophobic motifs in VPS37A identified above, promotes ESCRT-I targeting for autophagosome completion and the subsequent formation of a functional autolysosome.

### VPS37A UEVL domain remodels liposomes to high-order structures

Interestingly, VPS37A^21-148, 4A^ did not completely abolish its membrane binding (Fig. 3b and 3c). This prompted us to investigate additional membrane interactions of VPS37A^21-148, 4A^. Extended Data Fig. 8a shows an overlay of 2D ^1^H-^15^N correlation spectra of VPS37A^21-148, 4A^ in the presence and absence of bicelles. Residues having small chemical shift perturbations are distributed throughout the UEVL domain (Extended Data Fig. 8b). Furthermore, the intensities of 1D ^1^H-^15^N correlation spectra of VPS37A^21-148, 4A^ in bicelles decreased over time even though the sample remained visually clear and the protein alone was stable in aqueous solution (Figure 5a). This observation suggests that VPS37A^21-148, 4A^ protein in the presence of bicelles may form high molecular weight aggregates that are above the size detectable by solution NMR. To test this idea, we used DLS to measure the sizes and size-distributions of liposomes alone or incubated with VPS37A^1-148^ and VPS37A^21-148, 4A^. Changes in these parameters reflect the extent of membrane perturbations resulting from protein interactions. As shown in Fig. 5b-d, sonicated liposomes were stable, and few changes were detected over several days of observation. However, when VPS37A^1-148^ or VPS37A^21-148, 4A^ were added, profound changes in liposome sizes were rapidly detected (at 10 minutes) and the measured liposome size increased with increasing incubation times. Furthermore, the magnitude of these changes was dependent on protein to lipid ratios and the presence of the two hydrophobic motifs identified above; DLS measurements indicated the production of larger vesicle sizes with VPS37A^1-148^ than with VPS37A^21-148, 4A^. By comparison, we observed much smaller changes in liposome sizes when VPS37A^1-148^ or VPS37A^21-148, 4A^ was added to extruded liposomes that have radii larger than 50 nm or sonicated liposomes without PG (Extended Data Fig. 9). This observation is consistent with a model where the VPS37A UEVL domain is primarily responsible for inducing highly curved liposomes to form higher-order structures while the two conserved hydrophobic motifs promote their formation in a curvature-sensitive manner.

**Fig. 5:**
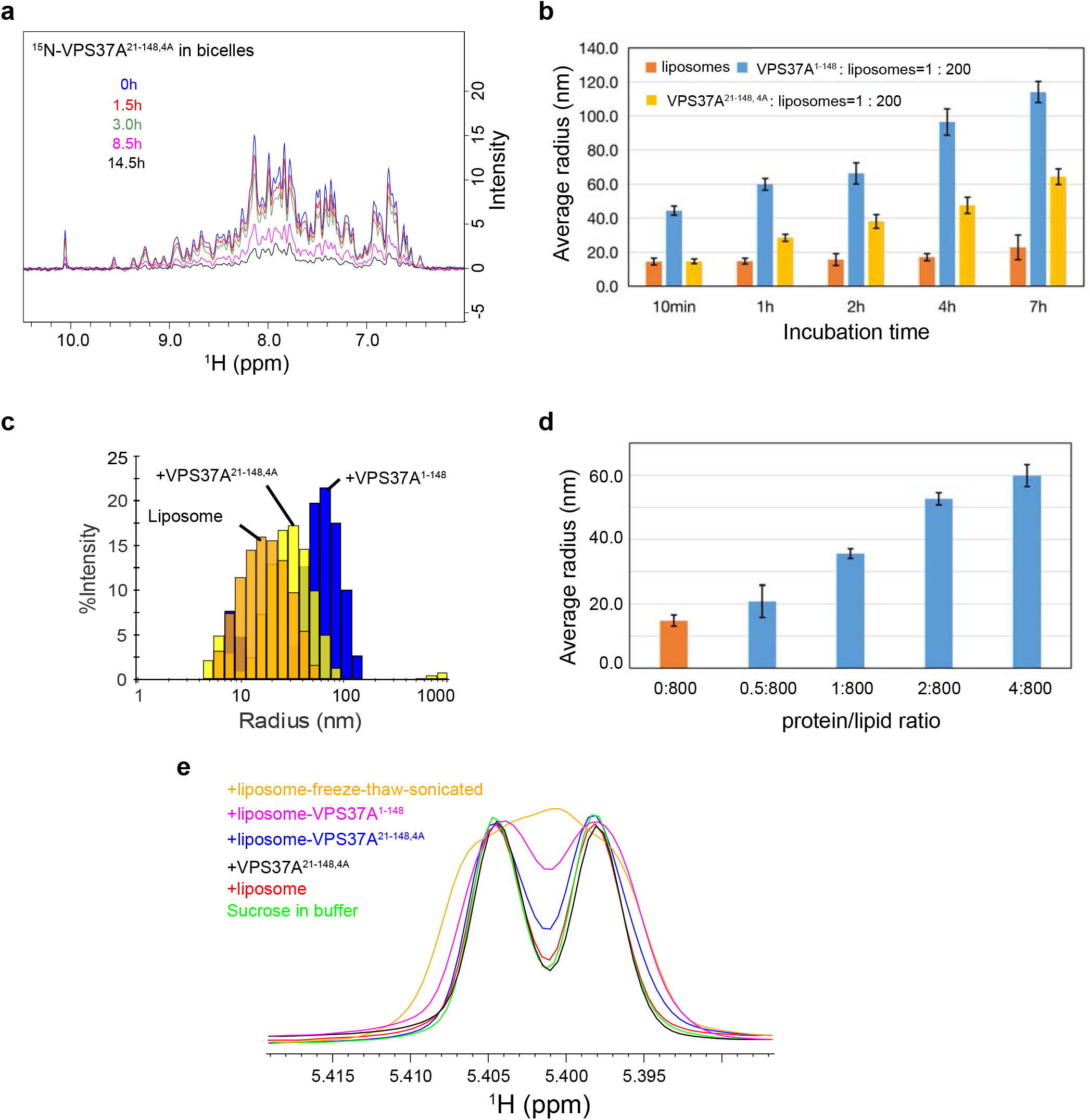
The VPS37A N-terminal induces formation of membrane high-order structures. a. 1D ^15^N-edited proton spectra of ^15^N-labeled VPS37A^21-148, 4A^ (100 μM) recorded at 15 ^o^C in 12% (w/v) bicelles (DMPC:DMPG:DHPC=8:2:20, q=0.5), 25 mM HEPES, pH 7.0, and 150 mM NaCl. b. Average radius of sonicated liposomes alone, or after mixing with VPS37A^21-148, 4A^ or VPS37A^1-148^ at different incubation times measured by DLS. 4 μM proteins were incubated with 800 μM sonicated liposomes (POPC:DOPG:DOPE=3:2:5) in 50 mM HEPES, pH 7.5, and 150 mM NaCl. c. Selected DLS profiles for sonicated liposomes (POPC:DOPG:DOPE=3:2:5) alone (orange), or with VPS37A^21-148, 4A^ (yellow), VPS37A^1-148^(blue) after 2 hrs incubation. d. Average radius of sonicated liposomes alone, or after mixing with VPS37A^1-148^ at different protein to lipid ratios measured by DLS after 1 hr incubation. e. Liposome leakage assay using NMR paramagnetic relaxation enhancement. 1D ^1^H spectra of 2 mM sucrose in buffer (green), with sonicated liposomes (red), with VPS37A^21-148, 4A^ (black), with liposomes and VPS37A^21-148, 4A^ (blue), with liposomes and VPS37A^21-148, 4A^ (magenta), and with disrupted liposomes (orange). Sonicated liposomes encapsulated with Mn^2+^ consist of POPC:DOPG:DOPE in a molar ratio of 3:2:5. Protein concentration is 2 μM and lipid concentration is 800 μM.

To determine if the formation of liposome high-order structures results in membrane leakage, we performed an NMR-based liposome-leakage assay using the paramagnetic relaxation enhancement (PRE) effect of Mn^2+ 20^. In this experiment, sucrose was mixed with sonicated liposomes encapsulated with Mn^2+^, and the resonance of its anomeric proton at ~5.4 ppm was monitored in the presence and absence of proteins using 1D ^1^H NMR. As shown in Fig. 5e, a minimal change in resonance linewidth was observed with the addition of Mn^2+^ encapsulated liposomes since the anomeric proton of sucrose experiences little PRE effect from the encapsulated metals. When VPS37A^21-148, 4A^ was mixed with Mn^2+^-encapsulated liposomes, some small changes in resonance linewidth were detected but these changes were smaller than those observed by the addition of disrupted liposomes. This suggests that the formation of high-ordered structures induced by VPS37A^21-148, 4A^ cause little liposome leakages and is likely a liposome tethering process. On the other hand, when VPS37A^1-148^ was mixed with Mn^2+^-encapsulated liposomes, moderate changes in resonance linewidth were detected due to some liposome leakages.

To directly visualize the effect of VPS37A^1-148^ binding on liposomes, we performed negative staining and EM experiment. As shown in Fig. 6, sonicated liposomes display relatively similar-sized vesicles with an average radius of about 20 nm. In contrast, following incubation with VPS37A^1-148^ this liposome preparation clearly resulted in the production of enlarged spherical vesicles and the formation of large rosette-like structures. We propose that the latter structures could be formed by clustering of multiple vesicles. Together, our results indicate that VPS37A^1-148^ efficiently remodels highly curved liposomes to form high-order structures *in vitro*.

**Fig. 6:**
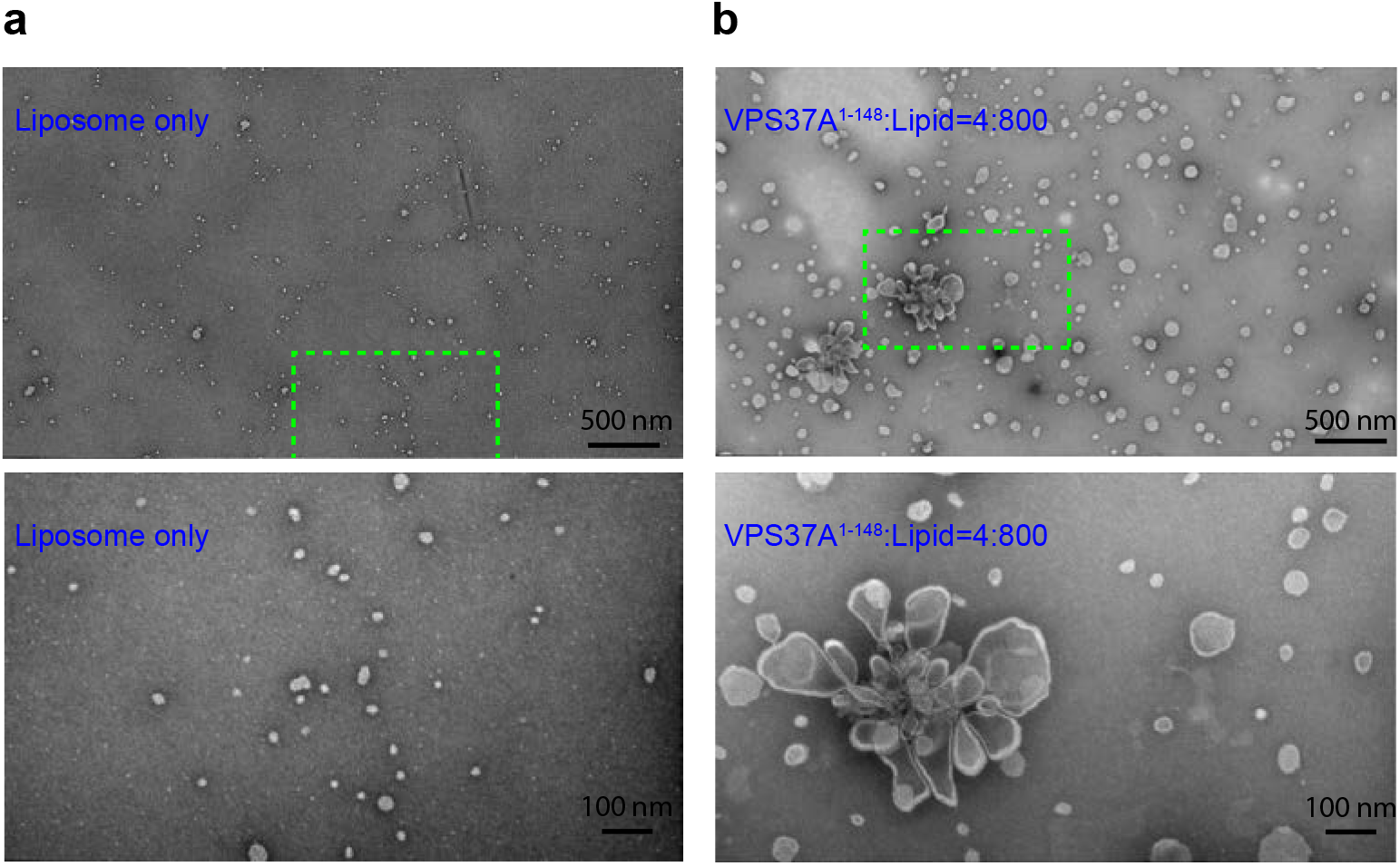
VPS37A^1-148^ clusters liposomes to high-order structures *in vitro*. Micrographs of negatively stained liposomes (POPC:DOPG=8:2) in the absence (a) and presence (b) of 0.8 μM VPS37A^1-148^ in 50 mM HEPES, pH 7.5, and 150 mM NaCl for 1 hr. Lipid concentration is 160 μM. Bottom panel shows zoomed in regions (green box) of top panel.

### VPS37A UEVL domain does not interact with mono-ubiquitin (Ub)

The UEV fold is present in a family of proteins structurally homologous to the canonical E2 ubiquitin-conjugating enzymes but lacks the catalytic cysteine residue^21^. Proteins containing the UEV fold are known to interact with Ub. For instance, TSG101 UEV binds to Ub. This interaction has been implicated in sorting of ubiquitinated cargos into the internal vesicles of multivesicular bodies^12, 13^; Mms2 (a UEV domain protein) forms a complex with Ub and E2 ligase Ubc13 for the assembly of polyubiquitin chains^22, 23^. Therefore, we examined the binding of the VPS37A UEVL domain to Ub using NMR chemical shift perturbation. 2D ^15^N-^1^H correlation spectra in the absence or presence of unlabeled Ub are superimposed in Extended Data Fig. 10a and show that few chemical shift perturbations were detected, demonstrating that VPS37A PUEV does not interact with Ub. By comparison, small but distinct spectral shifts for resonances from V43, D46, and G47 were observed in 2D ^15^N-^1^H correlation spectra of TSG101 UEV with and without unlabeled Ub as reported previously^12^ (Extended Data Fig. 10b). These residues have been shown to bind Ub from the so-called β-tongue of TSG101 UEV (Extended Data Fig. 10c). Extended Data Fig. 4 shows the secondary structure alignment and structure-based sequence alignment of VPS37A UEVL and TSG101 UEV. The ubiquitin-binding region in the TSG101 UEV (yellow regions in Extended Data Fig. 10c and 4a, and red boxed residues in Extended Data Fig. 4b) is absent in the VPS37A UEVL sequence providing an explanation for the experimental observation that VPS37A UEVL does not bind to Ub.

## Discussion

Autophagosome formation is a complex and dynamic membrane-remodeling process. It is primarily carried out by Atg proteins but factors that are responsible for phagophore closure have only recently been identified. Using a novel assay to differentiate between phagophores, autophagosomes, and autolysosomes, we presented the first experimental evidence that several components of the ESCRT machinery, such as VPS37A and CHMP2A, participate in phagophore closure in mammalian cells^5, 6^. Subsequent reports from other labs also confirmed that ESCRT is required for closing the phagophore in budding yeast and mitophagy^19, 24^. Together, these studies have established that ESCRT proteins mediate phagophore closure during the biogenesis of autophagosomes. In the present study, we have determined the structure of the residues 1 to 148 in the VPS37A and confirmed that it contains a UEVL domain. Although its core structure is similar to that of other UEV proteins, the VPS37A UEVL does not bind Ub as it lacks the ubiquitin interacting region of TSG101 UEV. In addition, we have discovered that the VPS37A N-terminal domain remodels highly curved liposomes to high-order molecular structures. This process is facilitated by two hydrophobic motifs in disordered regions that selectively interact with highly curved membranes and promote liposome clustering. The functional significance of VPS37A’s curvature-sensitive membrane interaction is demonstrated by *in vivo* mutagenesis studies; mutations in these two motifs compromise its phagophore localization and autophagic flux.

Membrane geometry has recently emerged as an essential component of the microenvironments in which membrane fusion and fission, protein localization, trafficking, and signaling occur. The curvature can regulate the activity and determine the subcellular localization of some proteins. For instance, human ArfGAP1 and Atg3 proteins display enhanced enzymatic activity when interacting with small lipid vesicles that are highly curved, but show little to no activity on large and less curved lipid vesicles ^17, 25^. Bacterial DivIVA and SpoVM proteins recognize membranes with negative or small positive curvatures for their intracellular localizations, respectively^26–28^. These proteins insert an amphipathic helix or a hydrophobic loop into one leaflet of the bilayer to sense membrane curvatures, which is one of the most common curvature recognition mechanisms. Membrane curve-sensitive binding is a recurring theme for autophagy-related proteins, including Atg1, Atg3, Atg13, Atg14L, and Atg16L1^29^. Previously, the selective binding of Atg3 to membranes with strong curvatures has been implicated in targeting the protein to the tip of phagophore during expansion^17^. Here we discovered two hydrophobic motifs, ^3^WLFP and ^137^FPYL, that mediate the preferential interaction of residues 1 to 148 of VPS37A with small and highly curved liposomes. Both sequences contain bulky hydrophobic sidechains and are effective for membrane insertion^16, 30^. Deleting these residues or substituting them with Ala greatly reduces VPS37A N-terminal binding to liposomes *in vitro* and compromises its localization to the phagophore and disrupts autophagic flux *in vivo*. Therefore, even though other interactions between VPS37A and phagophore factors remain to be determined, we propose that membrane curvature-sensitive interactions via two conserved hydrophobic motifs lead or contribute to the recruitment of VPS37A to the phagophore since highly curved membranes are a hallmark of its leading edge.

Human VPS37 has four homologues, VPS37A-D. While each contains a Mod(s) domain that interacts with other components of ESCRT-I to form the core structure, VPS37A has a unique large N-terminal domain (Extended Data Fig. 2). In a previous genome-wide screening we found that VPS37A, but none of its homologues, is involved in phagophore closure, and the N-terminal domain is indispensable for targeting VPS37A to the phagophore^5^. Despite intense efforts we have not been able to identify protein molecules on the phagophore that interact with and recruit VPS37A. However, the membrane curvature-selective interaction of the N-terminal domain uncovered in this study provides a model for how the phagophore’s distinct and highly curved rim may function as a geometric cue to recruit VPS37A. By comparison, the ESCRT-1 complex is recruited to nearly “flat” membranes of endosomes during initiation via interactions with ESCRT-0 and ubiquitinated cargos in the MVB pathway. Thus, while VPS37A-D has redundant function in the MVB pathway, differences in membrane structures may explain why deletion of the VPS37A N-terminal domain disrupts its phagophore recruitment but has little effect on its role in the degradation of the epidermal growth factor receptor.

The observation that the VPS37A N-terminal induces liposome clustering and leakage *in vitro* indicate that beyond curvature sensing it may perturb bilayer structures^31,32^. While biological implications of this observation remain to be determined, several Atg proteins including Atg2A, Atg5, Atg8, Atg14, and Atg17 reportedly mediate membrane tethering^33–37^. Interestingly, membrane tethering by GATE-16 and LC3B proteins, human homologs of yeast Atg8, is curvature sensitive: GATE-16 successfully tethers large lipid vesicles (e.g. 100 and 200 nm in radius), while LC3 preferentially tethers liposomes with a radius smaller than 25 nm^38^. Here we demonstrate that the VPS37A N-terminal has an intrinsic ability to remodel negatively charged and highly curved liposomes. Since the leading edge of the cuplike phagophore shares these membrane characteristics, it is tempting to speculate that the VPS37A N-terminal may facilitate ESCRT-III/VPS4 dependent phagophore closure by destabilizing lipid bilayers at the scission site. Future studies are required to investigate this hypothesis and the underlying molecular mechanisms.

## Materials and Methods

### Reagents

The following antibodies were used for immunoblotting: β-ACTIN (Sigma-Aldrich, A5441, 1:10,000); GFP (Cell Signaling, 2956, 1:1,000); MAP1LC3B (Novus, NB100-2220, 1:3,000); p62 (American Research Products, 03-GP62-C, 1:4,000). ON-TARGETplus SMART Pool Non-targeting (D-001810-10) and CHMP2A (L-020247-01) siRNAs were obtained from GE Healthcare Dharmacon. pCDH1-CMV-HA-VPS28(WT)-SV40-hygro and pCDH1-CMV-HA-VPS28(T2[K54D, K58D, D59A])-SV40-hygro were generated using Gibson Assembly. All other reagents were obtained from the following sources: Bafilomycin A1 (LC Laboratories, B-1080); Hoechst 33342 (Invitrogen, NucBlue, R37605); Membrane-impermeable HaloTag Ligand (MIL) (Promega, Alexa Fluor 488-conjugated, G1001); Membrane-permeable HaloTag Ligand (MPL) (Promega, tetramethylrhodamine-conjugated, G8251); normal goat serum (Sigma-Aldrich, G9023); Nucleofector Kit V (Lonza, VCA-1003); paraformaldehyde (Electron Microscopy Sciences, 15710); XF Plasma Membrane Permeabilizer (XF-PMP) (Seahorse Bioscience, 102504-100).

To construct pCDH1-CMV-GFP-VPS37A^Δ20,4A^-EF1-Puro, pCDH1-CMV-GFP-VPS37A^4A^ was first generated by Gibson Assembly using pCDH1-CMV-GFP-FL(VPS37A WT)-EF1-Puro^5^ and the following sets of primers: VPS37A^4A^-F1/R1; VPS37A^4A^-F2/R2. The resultant plasmid and the primer set VPS37A-dN20-F/R were used to generate VPS37A^Δ20, 4A^ cDNA, which was then subcloned into the XhoI/BamHI site of pCDH1-CMV-GFP. The UEV^TSG101-^VPS37A expression construct (pCDH1-CMV-GFP-TSG101^1-145^-VPS37A^150-397^) was generated by Gibson Assembly using pLNCX2-mEGFP-TSG101 (a gift from Dr. Sanford Simon, Addgene plasmid # 116925), pCDH1-CMV-GFP-FL(VPS37A WT)-EF1-Puro, and the following primer sets: TSG101 UEV-F/R; VPS37A^150-397^-F/R. The primer sequences are listed in Supplementary Table 2.

### Cell culture and viral transduction

U-2 OS and 293T/17 (CRL-11268) cells obtained from American Type Culture Collection were maintained in McCoy’s 5A Medium supplemented with 10% fetal bovine serum (FBS) or Dulbecco’s Modification of Eagle’s Medium (DMEM) supplemented with 10% FBS, respectively. VPS37A knockout U-2 OS cells stably expressing GFP and GFP-VPS37A (WT) were generated as described previously^5^. Lentiviral production and transduction were conducted using the Invitrogen ViraPower Lentiviral Expression System as described previously^5^.

### Immunoblotting, autophagic flux assay and confocal microscopy

Total cell lysates were prepared in radio-immunoprecipitation assay buffer (150 mM NaCl, 10 mM Tris-HCl, pH 7.4, 0.1% SDS, 1% Triton X-100, 1% Deoxycholate, 5 mM EDTA, pH 8.0) containing protease inhibitors and subjected to SDS-PAGE followed by immunoblotting with indicated antibodies as described previously ^6^. For autophagic flux assay, cells were rinsed three times with Dulbecco’s Phosphate Buffered Saline and incubated with amino acid-free DMEM in the presence or absence of 100 nM BafA1 as described previously^5^. The signal intensities were quantified using the Image Studio version 5 software (LI-COR Biotechnology) and starvation-induced autophagic flux was calculated as described previously^5^. For confocal microscopy, cells grown on Lab-TekII Chambered Coverglass, Chamber Slide (Nunc, 154941) were fixed in 4% paraformaldehyde-PBS at room temperature (RT) for 10 min. Fluorescence images were obtained using a Leica AOBS SP8 laser-scanning confocal microscope (63x oil-immersion [1.2 numerical aperture] lens) with the highly sensitive HyD detectors and the Leica image acquisition software LAX, deconvolved using Huygens deconvolution software (Scientific Volume Imaging), and analyzed using Imaris software (Bitplane) and Volocity software (PerkinElmer) without gamma adjustment.

### Statistical analyses

Statistical significance was determined using Graph Pad Prism 7.0. The threshold for statistical significance for each test was set at 95% confidence (p<0.05).

### Protein expression and purification

DNA sequences encoding residues 1 to 148 of the VPS37A N-terminal (VPS37A^1-148^) and residues 1 to 145 of the TSG101 UEV domain were separately subcloned into the pET28a expression vector at the BamHI/XhoI site with a His6-tag, a T7-tag and a thrombin cleavage site between the T7-tag and N-termini. All VPS37A^1-148^ mutants were generated using the Q5 Site-Directed Mutagenesis Kit (New England Biolabs). The corresponding primers are listed in Supplementary Table 2 and all constructs were verified by sequencing. Plasmids were then transformed into chemically competent Rosetta™(DE3) pLysS cells for expression. Typically, a single colony was selected to grow in a small volume of LB medium overnight at 37 °C as a starter culture and then inoculated into a large volume of LB medium (for unlabeled proteins) or M9 medium supplemented with D-glucose (or 3 g/L D-glucose-^13^C6) and ^15^NH4Cl (1 g/L) for ^15^N (or ^15^N/^13^C) labeled samples at 37 °C. After OD_600_ reached 0.6 to 0.8, the temperature was lowered to 25 °C and cells were induced with 0.5 mM IPTG for ~16 hrs. Cells were harvested by centrifugation and the pellets were stored at –80 °C until use.

Cell pellets for VPS37A^1-148^ or its mutants were homogenized with a lysis buffer of 20 mM phosphate, pH 7.5, 300 mM NaCl, 2 mM β-mercaptoethanol (BME), complete protease inhibitor cocktail (Roche), and 0.1% triton X-100 (Alfa Aesar) and were lysed by sonication on ice with 2 s on and 7 s off intervals for 18 mins total duration or by microfluidics. Cell debris was removed by centrifugation (Sorvall RC5B Plus Refrigerated Centrifuge) at 26,900 *×g* at 10 °C for 30 mins. Supernatants were collected and loaded onto a Ni-NTA column (HisTrap HP). The column was washed with PBS buffers of 20 mM phosphate, pH 7.5, 300 mM NaCl, 2 mM BME without and with 20 mM imidazole, and then eluted with a PBS buffer containing 500 mM imidazole. The elution of VPS37A^1-148^ or its mutants from the Ni-NTA column was concentrated and exchanged into a buffer containing 50 mM HEPES, pH 7.5, 150 mM NaCl, and 2 mM BME, and followed by the addition of 0.1% (v/v) TWEEN20 (Fisher Biotech) and 50 units thrombin (EMD Millipore) to remove the T7-tag and His6-tag for an overnight agitation at 4 °C. The solution was then subjected to a Ni-NTA column and the flow-through was collected and further purified by size-exclusion chromatography using a S200 column (HiLoad 16/60 Superdex 200) with a running buffer of 50 mM HEPES, pH 7.5, 1 M NaCl, and 1mM DTT. Purified VPS37A^1-148^ or mutant proteins were exchanged into a buffer of 20 mM PBS, pH 6.5 (or 50 mM HEPES, pH 6.5 to 7.5), 150 mM NaCl. Protein concentration was assayed using a Nanodrop (Thermo Scientific, Waltham, MA). TSG101 UEV was purified with a similar protocol, and the purified protein was exchanged into a buffer of 50 mM HEPES, pH 6.8, 150 mM NaCl, and 2 mM TCEP.

### Liposome flotation assays

Liposomes were prepared in H_2_O by sonication or extrusion with membrane pore sizes of 50, 100, and 200 nm as described previously^18^. Proteins (typically 2 to 4 μM) were incubated with 800 μM liposomes in buffer A containing 50 mM HEPES, pH 7.5, and 150 mM NaCl (total volume of 300 μL) at room temperature for 1 hr. The 300 μL mixtures were adjusted to a 40% OptiPrep Density gradient medium by mixing with 200 μL 100% (w/v) OptiPrep Density gradient medium (Sigma). The mixture was transferred to a 3.2 mL, Open-Top Thickwall Polypropylene Tube (13 x 56) mm at the bottom, followed by a 416.7 μL middle layer of 30% OptiPrep Density gradient medium and an 83.3 μL top layer of 0% OptiPrep Density gradient medium which were prepared separately in buffer A. Each tube was then subjected to centrifugation at 200,000 ×*g* in a sw55Ti rotor (Beckman) at 10 °C for 4 h. 125 μL samples from the top, middle and bottom of the gradient were collected and 15 μL of each were mixed with 4 μL 4x loading buffer and heated for 3 mins before SDS-PAGE gel electrophoresis (SurePAGE 10% Bis-Tris, GenScript).

### Dynamic Light Scattering Measurement

VPS37A^1-148^ or its mutants (0.5 to 4 μM) was incubated with 800 μM liposomes in buffer A (total volume of 30 μL) at room temperature and each mixture was measured three times at each time point by DLS (WYATT, DynaPro NanoStar) at 25 °C. Data was processed and analyzed using DYNAMICS V7 software.

### Liposome leakage assay by Mn^2+^ PRE NMR

POPC, DOPG and DOPE lipids with a molar ratio of 3:2:5 (POPC:DOPG:DOPE) were solubilized in chloroform and dried to a thin film by SpeedVac, followed by lyophilization overnight. Lipids were rehydrated in a 1 mL buffer A containing 10 mM MnCl_2_ for 1 hr at 42 °C with vortex-freeze-thaw every 20 mins, and then sonicated for 5×15-min intervals until clear. The prepared liposomes (4 mM lipids, 1 mL) were dialyzed once against a 150 mL buffer A containing 1 mM EDTA once and then twice against buffer A (150 mL each time). A 2 mM sucrose stock solution was prepared in a D_2_O buffer of 50 mM HEPES, pH 7.1 (measured by pH meter, corresponding to pH 7.5 in H_2_O buffer), and 150 mM NaCl. Liposomes in 4 mM stock were added to the sucrose solution to a final concentration of 800 μM, and proteins (VPS37A^21-148, 4A^ or VPS37A^1-148^) were added to a final concentration of 2 μM. 1D ^1^H NMR spectra were acquired at 25 °C.

### Transmission electron microscopy

4 μM VPS37A^1-148^ were incubated with 800 μM liposomes in buffer A (total volume of 60 μL) for 1 hr at room temperature and then diluted in H_2_O to a final volume of 300 μL. Negatively stained samples were prepared by applying a 10 μL sample to Formvar-coated 400 mesh copper grids for 2 mins, blotting excess sample, staining for 2 mins in 1% (w/v) uranyl acetate, and a final blotting of excess stain. The grid was allowed to dry and was examined the same day. Micrographs were taken at a magnification of 12,000 or 40,000 with a Jeol JEM 1400 transmission electron microscope (JEOL USA Inc.) operating at 60 kV.

### NMR spectroscopy and structure determination

Typical NMR samples contained 0.05 to 0.5 mM labeled proteins in a buffer of 20 mM PBS, pH 6.5 (or at specified buffer conditions) containing 150 mM NaCl and 0.02% NaN3. For residual dipolar coupling measurements, 140 μM ^15^N-labeled VPS37A^21-148^ was dissolved in 50 mM HEPES, pH 6.5, and 250 mM NaCl containing 6.5 mg/mL phage Pf1 (ASLA BIOTECH). For bicelle samples, 50 μM ^15^N-labeled VPS37A^1-148^ were mixed with a final concentration of 12% (w/v) bicelles (DMPC:DMPG:DHPC = 4:1:10, molar ratio, q=0.5) in 25 mM HEPES, pH 7.0, 150 mM NaCl. All NMR experiments were conducted at 25 °C except the one in bicelles that was collected at 15°C (since the sample precipitates after NMR collection at 25 °C). All NMR data were acquired on a Bruker 600 MHz spectrometer except the NOESY data that were acquired on a Bruker 850 MHz spectrometer. Both instruments are equipped with cryoprobes. The data were processed using NMRPipe and analyzed using NMRView. Backbone resonance assignments were carried out using triple resonance HNCO, HN(CA)CO, HNCA, HN(CO)CA, HNCACB, and HN(CO)CACB experiments. Aliphatic and aromatic sidechain resonance assignments were obtained from 3D ^13^C-edited HCCHTOCSY, ^13^C-edited NOESY, ^15^N-edited NOESY and HBHA(CBCACO)NH, 2D Hb(CbCgCd)Hd and Hb(CbCgCdCe)He spectra. The 3D ^13^C-NOESY, ^15^N-NOESY experiments were collected with 150 ms mixing time. ^15^N-^1^H RDCs were measured using the IPAP-HSQC experiment.

Torsion angle constraints were derived from C_α_, C_β_, N, and C’ chemical shifts using TALOS+. NOE assignments were obtained by a combined manual and automated analysis with CYANA 3.0. The structures were calculated with XPLOR-NIH 3.3. PSVS was used to analyze the 10 lowest-energy structures.

## Data Availability

NMR resonance assignments have been deposited with the BMRB with accession number #51558 for VPS37A^1-148^ and #31039 for VPS37A^21-148^. NMR structure has been deposited with the Protein Data Bank with accession ID 8E22. All other data that support the findings of this study are available from the corresponding authors upon reasonable request.

## Acknowledgements

We thank Dr. Han Chen for his help with TEM experiments. We are thankful for financial support from the National Institutes of Health NIGMS (R01 GM127730, R01 GM127954, R01 CA222349) and Four Diamonds to FT and JMF. The NMR core, Advanced Light Microscopy core (RRID:SCR_022526), and TEM Core (RRID: SCR_021200) services and instruments used in this project were funded, in part, by the Pennsylvania State University College of Medicine via the Office of the Vice Dean of Research and Graduate Students and the Pennsylvania Department of Health using Tobacco Settlement Funds (CURE). The content is solely the responsibility of the authors and does not necessarily represent the official views of the University or College of Medicine. The Pennsylvania Department of Health specifically disclaims responsibility for any analyses, interpretations, or conclusions.

## Author contributions

Y.S.Y., X.W.L., G.F.W., X.M.L., and F.T. designed and performed the experiments, analyzed the data, and drafted figures and the paper. M.C.B. and J.M.F. helped with VPS37A purification, data analysis, and manuscript preparation. F.T., Y.T., Y.S.Y., and H.G.W. conceived, planned, and supervised this study. F.T., Y.T., and Y.S.Y. wrote the final paper with feedback from all authors.

