## Supplemental Information for "Curvature Sensing and Membrane Remodeling of the VPS37A N-terminal Domain during Autophagy"

**Supplementary Table 1 | NMR constraints and structure statistics for VPS37A$^{21\text{-}148}$**
(NOE, Nuclear Overhauser Effect; RMSD, root-mean-square deviation)

| | *XPLOR-NIH* |
|---|---|
| ***NMR constraints*** | |
| *NOE distances* | 2385 |
| intra-residue | 538 |
| sequential (\|*i-j*\| = 1) | 666 |
| medium range (2 ≤ \|*i-j*\| ≤ 4) | 417 |
| long range (\|*i-j*\| ≥ 5) | 764 |
| *Dihedral angles* | |
| phi | 97 |
| psi | 97 |
| *Residual dipolar couplings ($D_{NH}$)* | 74 |
| ***Cis-prolines*** | P74, P84 |
| ***Structure statistics*** | |
| *Ensemble RMSD** | |
| backbone heavy atoms (Å) | 0.3 |
| all heavy atoms (Å) | 0.9 |
| *Ramachandran analysis*** | |
| most favored | 98.8% |
| additionally allowed | 1.2% |
| generally allowed | 0.0% |
| disallowed | 0.0% |
| *Violations (mean ± standard deviation)* | |
| distance constraints (Å) | 0.066 ± 0.001 |
| dihedral angle constraints (°) | 0.667 ± 0.093 |
| Residual dipolar couplings (Hz) | 0.925 ± 0.060 |
| *Deviations from idealized geometry* | |
| bond lengths (Å) | 0.004 ± 0.000 |
| bond angles (°) | 0.611 ± 0.006 |
| improper (°) | 0.504 ± 0.010 |

* Evaluated for secondary structure elements: 24-39, 44-45, 51-58, 61-68, 79-83, 86-87, 101-104, 112-125, 129-130.

** Evaluated by Procheck for secondary structure elements: 24-39, 44-45, 51-58, 61-68, 79-83, 86-87, 101-104, 112-125, 129-130.

**Supplementary Table 2 | Primers (F, Forward; R, Reverse) used in this study.**

| **Primers ID for in vitro experiments** | Sequences from 5' to 3' |
|---|---|
| VPS37A$_{1-148}$_F | AGTGCCTCGCGGATCCATGAGCTGGCTTTTTC |
| VPS37A$_{1-148}$_R | GGTGGTGGTGCTCGAGCTAAGACATCCCACTT |
| VPS37A$_{21-148}$_F | GGCCTCACCAGCCTCCAG |
| VPS37A$_{21-148}$_R | GGATCCGCGAGGCACTAG |
| VPS37A$_{1-148,4A}$/VPS37A$_{21-148,4A}$_F | GCTGCATACAGTAACCCAAGTGGG |
| VPS37A$_{1-148,4A}$/VPS37A$_{21-148,4A}$_R | TGCAGCTGCTGTTGAAGTAGGAGC |
| Tsg101 UEV_F1 | AGTGCCTCGCGGATCCATGGCGGTGTCGGAG AGC |
| Tsg101 UEV_F2 | GGTGGTGGTGCTCGAGCTACGGTCGTGAAAAT ACAGGTGG |
| **Primers ID for in cell experiments** | Sequences from 5' to 3' |
| VPS37A$^{4A}$-F1 | TACAAGTCCGGACTCAGATCTCGAGCTCAAAG CTGGCTTTTTC |
| VPS37A$^{4A}$-R1 | TGCTGTTGAAGTAGGAGCTAAAAC |
| VPS37A$^{4A}$-F2 | TAGCTCCTACTTCAACAGCAGCTGCAGCAGCA TACAGTAACCCAAGTGGG |
| VPS37A$^{4A}$-R2 | CGCAGATCCTTGCGGCCGCGGATCCCTATAGT GGAGCATG |
| VPS37A$_{21-397}$-F | TACAAGTCCGGACTCAGATCTCGAGCTGGCCT CACCAGCCTCCAG |
| VPS37A$_{21-397}$-R | CGCAGATCCTTGCGGCCGCGGATCCCTATAGT GGAGCATGAAATTGG |
| TSG101 UEV-F | TACAAGTCCGGACTCAGATCTCGAGCTCAAGC TTCGAATTC |
| TSG101 UEV-R | CGGTCGTGAAAATACAGG |
| VPS37A$_{150-397}$-F | CACCTGTATTTTCACGACCGTATGCTTCTCAGG GTTTTC |
| VPS37A$_{150-397}$-R | CGCAGATCCTTGCGGCCGCGGATCCCTATAGT GGAGCATG |

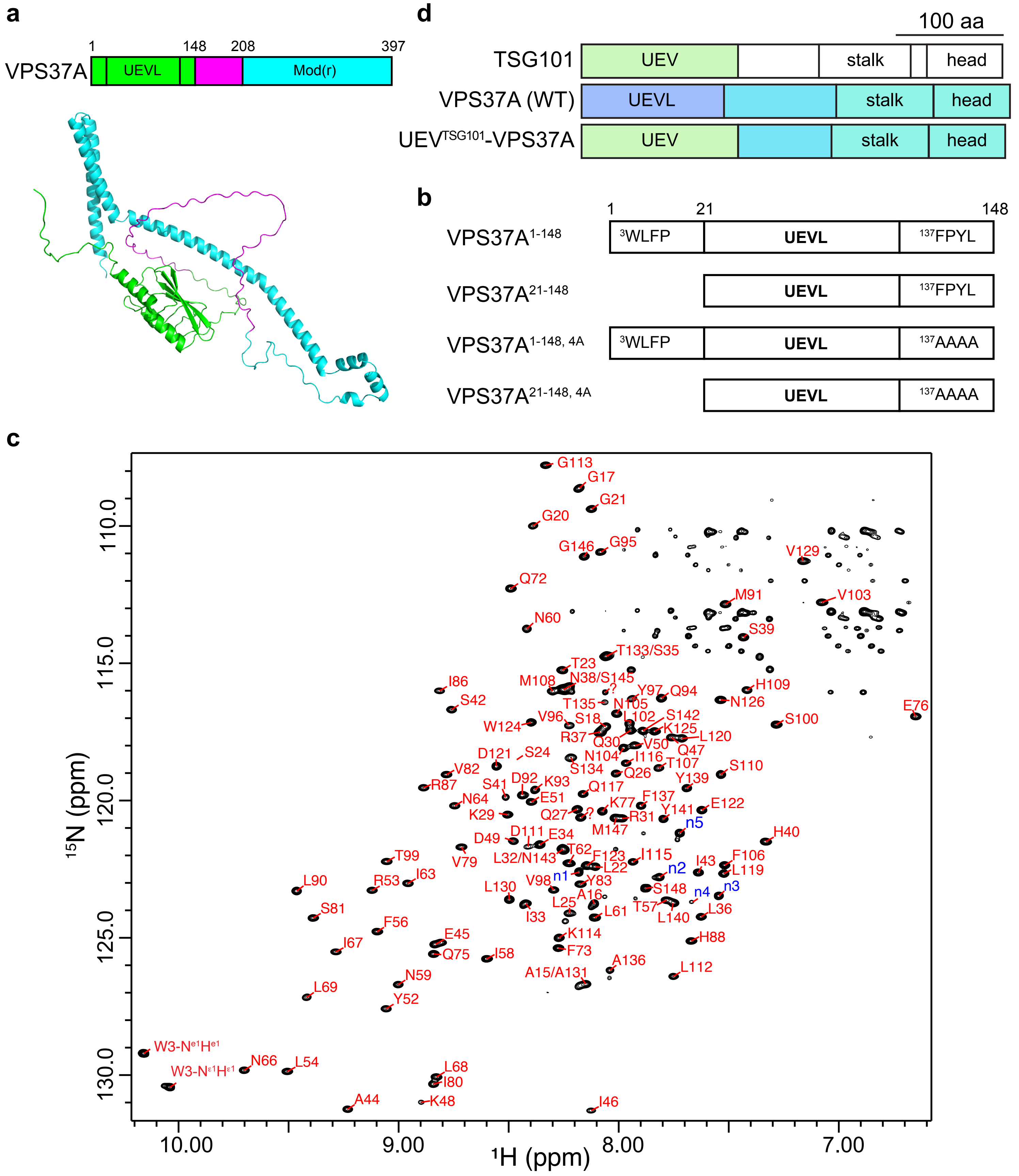


**Extended Data Fig. 1| VPS37A domain structure and constructs used in this study.**
**a,** Human VPS37A domain structure and its predicted AlphaFold structure. The N-terminal containing a UEVL domain is shown in green, while the C-terminal containing a modifier of rudimentary [Mod(r)] domain is shown in cyan. A long disordered stretch between the N- and C-termini is shown in magenta.
**b,** Diagrams of VPS37A N-terminal constructs for *in vitro* study.
**c,** $^{15}N$-$^{1}H$ TROSY spectrum of $^{15}N$, $^{13}C$-labeled VPS37A$^{1\text{-}148}$ with resonance assignments. The unassigned residues from N-terminal are indicated as n1 to n5. The spectrum was acquired on a Bruker 600 MHz spectrometer at 25 °C.
**d,** Diagrams of VPS37A and a VPS37A mutant for *in vivo* study.

```
VPS37A      MSWLFPLTKSASSSAAGSPGGLTSLQQQKQRLIESLRNSHSSIAEIQKDVEYRLPFTINN      60
VPS37D      ------------------------------------------------------------      0
VPS37B      ------------------------------------------------------------      0
VPS37C      ------------------------------------------------------------      0

VPS37A      LTININILLPPQFPQEKPVISVYPPIRHHLMDKQGVYVTSPLVNNFTMHSDLGKIIQSLL      120
VPS37D      ------------------------------------------------------------      0
VPS37B      ------------------------------------------------------------      0
VPS37C      ------------------------------------------------------------      0

VPS37A      DEFWKNPPVLAPTSTAFPYLYSNPSGMSPYASQGFPFLPPYPPQEANRSITSLSVADTVS      180
VPS37D      ------------------------------------------------------------      0
VPS37B      ------------------------------------------------------------      0
VPS37C      ------------------------------------------------------------      0

VPS37A      SSTTSHTTAKPAAPSFGVLSNLPLPIPTVDASIPTSQNGFGYKMPDVPDAFPELSELSVS      240
VPS37D      ---------------------------MYRA--------RAARAGPEPGSPGRFGILSTG      25
VPS37B      --------------------------------------------MAGAGSEARFAGLSLV      16
VPS37C      --------------------------------------------------METLKDKTLQ      10
                                                                 :   :

VPS37A      QLTDMNEQEEVLLEQFLTLPQLKQIITDKDDLVKSIEELARKNLLLEPSLEAKRQTVLDK      300
VPS37D      QLRDLLQDEPKLDRIVRLSRKFQGLQLEREACLASNYALAKENLALRPRLEMGRAALAIK      85
VPS37B      QLNELLEDEGQLTEMVQKMEETQNVQLNKEMTLASNRSLAEGNLLYQPQLDTLKARLTQK      76
VPS37C      ELEELQNDSEAIDQLALESPEVQDLQLEREMALATNRSLAERNLEFQGPLEISRSNLSDR      70
            :* :: ::.  : .      : : :  :::  : :   **. **  .  *:  :  :  :

VPS37A      YELLTQMKSTFEKKMQRQHELSESCSASALQARLKVAAHEAEEESDNIAEDFLEGKMEID      360
VPS37D      YQELREVAENCADKLQRLEESMHRWSPHCALGWLQAELEEAEQEAEEQMEQLLLGEQSLE      145
VPS37B      YQELQVLFEAYQIKKTKLDRQSSSASLETLLALLQAEGAKIEEDTENMAEKFLDGELPLD      136
VPS37C      YQELRKLVERCQEQKAKLEKFSSALQPGTLLDLLQVEGMKIEEESEAMAEKFLEGEVPLE      130
            *: *  : .    :  : ..     .       *:.   : *::::   *.:* *:  ::

VPS37A      DFLSSFMEKRTICHCRRAKEEKLQQAIAMHSQFHAPL-----------------------      397
VPS37D      AFLPAFQRGRALAHLRRTQAEKLQELLRRRERSAQPAPTSAADPPKSFPAAAVLPTGAAR      205
VPS37B      SFIDVYQSKRKLAHMRRVKIEKLQEMVLKGQRLPQALAPL--PPRL------------PE      182
VPS37C      TFLENFSSMRMLSHLRRVRVEKLQEVVRKPRASQELAGDA--PPPRPPPPVRPVPQGTPP      188
             *:  :   * :.* **.: ****: :

VPS37A      ------------------------------------------------------------      397
VPS37D      G---PPA-----------------------VPRSLP------------------------      215
VPS37B      LA---PT---APLPYPAPEASGPP----AVAPRRIPPPPPPVP------AGRLATPFTAA      226
VPS37C      VVEEQPQPPLAMPPYPLPYSPSPSLPVGPTAHGALPPAPFPVVSQPSFYSGPLGPTYPAA      248

VPS37A      ------------------------------------------------------------      397
VPS37D      -----------PLDSRPVPPLKGSPGCPLGPAPLL------------------SPRPS--      244
VPS37B      M-SSGQAVPYPGLQCPPLPPRVGLP------------------TQQGFSSQFVSPYPP--      265
VPS37C      QLGPRGAAGYSWSPQRSMPPRPGYPGTPMGASGPGYPLRGGRAPSPGYPQQ--SPYPATG      306

VPS37A      ----------------------------------------------------      397
VPS37D      ----------------QPEPPHR-----------------------------      251
VPS37B      ------------PLPQRPPP--------------------RLPPHQPGFILQ      285
VPS37C      GKPPYPIQPQLPSFPGQPQPSVPLQPPYPPGPAPPYGFPPPPGPAWPGY---      355
```

**Extended Data Fig. 2 | Sequence alignment of four mammalian VPS37 homologues.**
The VPS37A has a unique N-terminal domain.

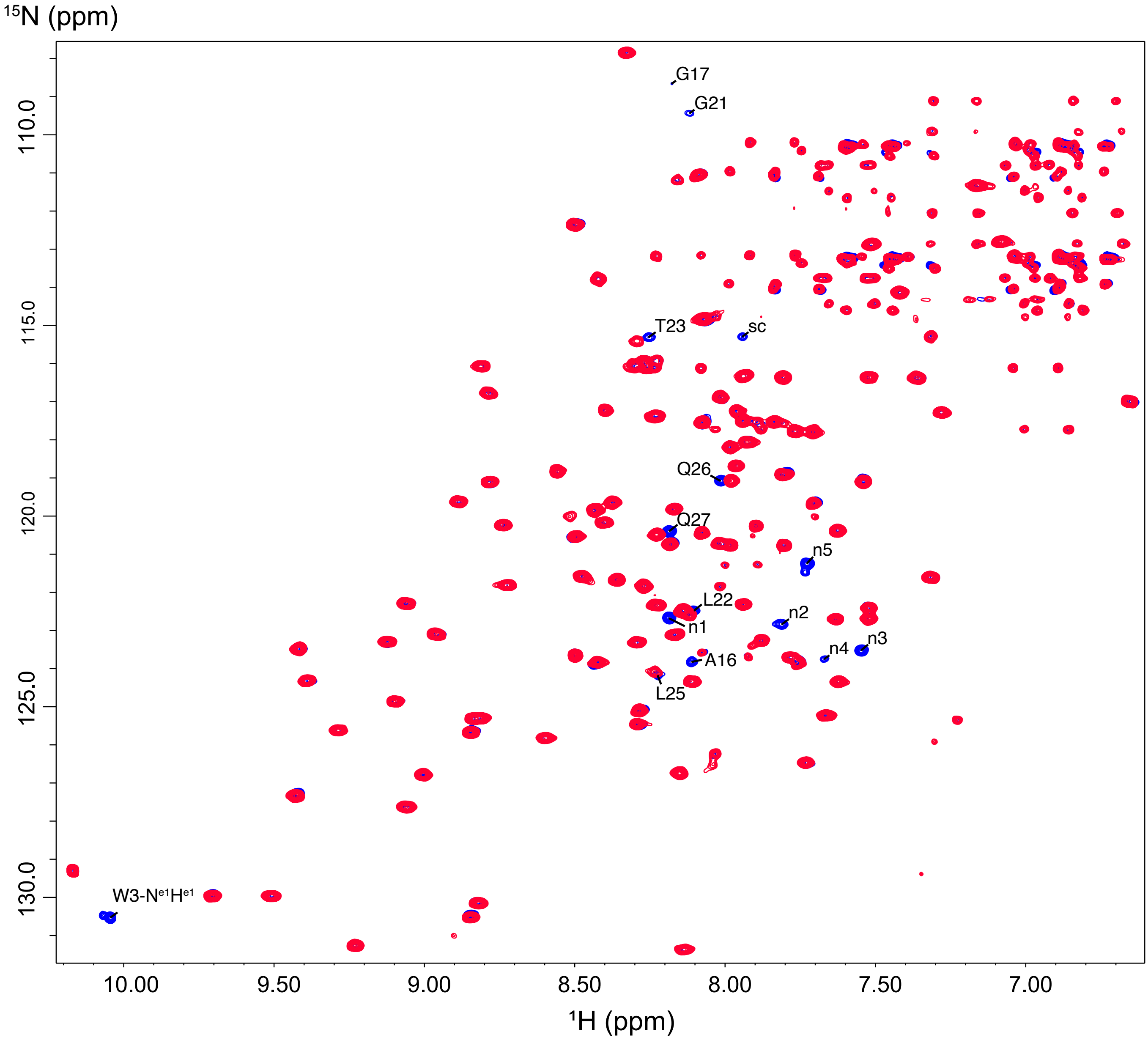


**Extended Data Fig. 3 | Deletion of N-terminal 20 residues does not perturb the UEVL domain structure.** Overlay of $^{15}$N-labeled VPS37A$^{1-148}$ (blue) and VPS37A$^{21-148}$ (red) TROSY spectra acquired on a Bruker 600 MHz spectrometer at 25 °C. Perturbed, unassigned (likely from the unassigned N-terminal), and sidechain resonances are indicated with their assignments, n1 to n5, and sc, respectively.

**a**

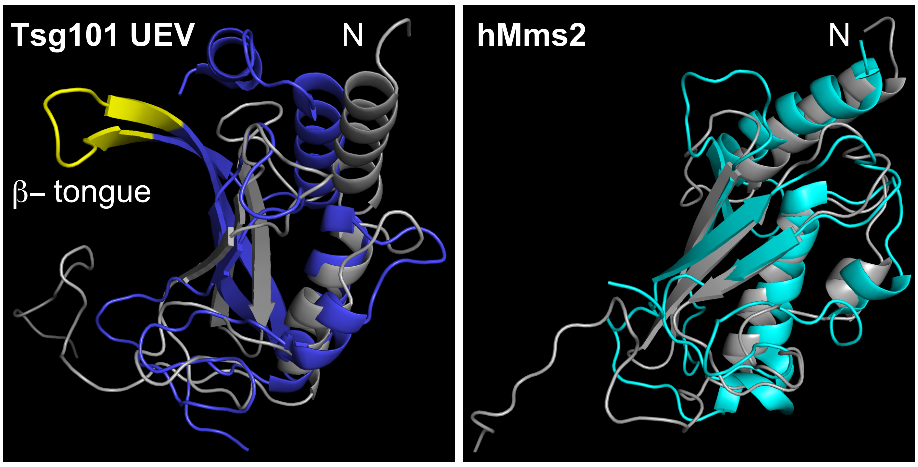


**b**

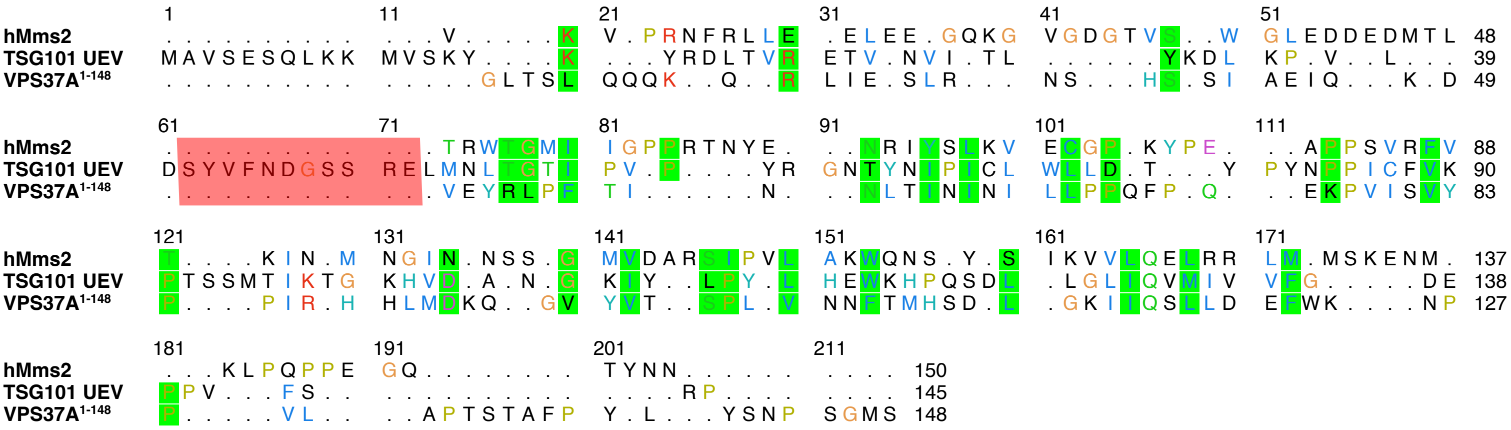


**Extended Data Fig. 4 | Structure and sequence alignments of VPS37A$^{1-148}$, human Mms2 (hMms2), and TSG101 UEV domain.**

**a,** Structural alignments of VPS37A N-terminl UEVL (gray) with TSG101 UEV (blue), and with hMms2 (cyan). β-tongue of TSG101 UEV is shown in yellow. N-terminus is indicated.

**b,** Structure-based sequence alignments of of VPS37A$^{1-148}$, hMms2, and TSG101 UEV. The β-tongue region from TSG101 UEV is highlighted in red.

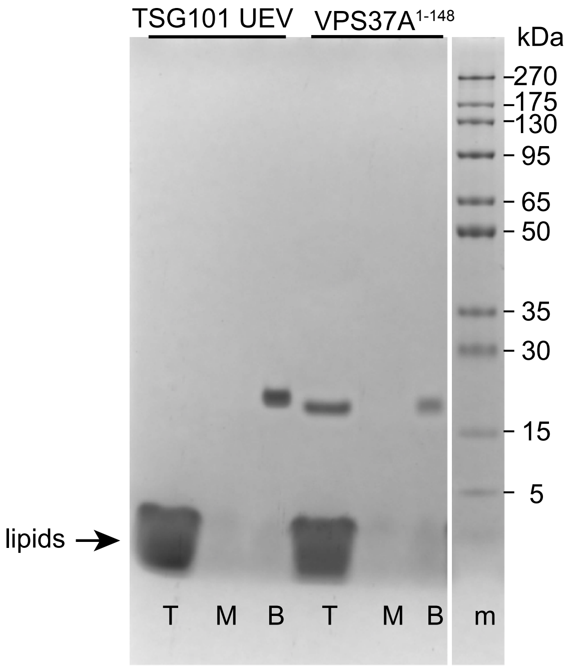


**Extended Data Fig. 5 | TSG101 UEV does not bind to sonicated liposomes.**

SDS-PAGE gel analysis of the co-flotation assays for TSG101 UEV and VPS37A$^{1\text{-}148}$ with sonicated liposomes (POPC:DOPG:DOPE=3:2:5, 800 μM) in a protein/lipid molar ratio of 1 to 400 in 50 mM HEPES, 150 mM NaCl, pH 7.5, 2 mM TCEP. T, M and B represent the top, middle and bottom layers, respectively, after density gradient ultracentrifugation. Protein marker is indicated by m. The arrow points to the lipid band.

```
VPS37A-Xenopus_laevis     MSWLFPSSKSNR-----PLQQLTSLQQQKQRQIESLKACHSSIIEIQKDVEYRLPFTVKN  55
VPS37A-Homo_sapiens       MSWLFPLTKSASSSAAGSPGGLTSLQQQKQRLIESLRNSHSSIAEIQKDVEYRLPFTINN  60
VPS37A-Mus_musculus       MSWLFPLAKSASSSAAGSPAGLTSLQQQKQRLIESLRNSHSSIAEIQKDVEYRLPFTVNN  60
                          ****** :**           ********** ****: .**** *************::*

VPS37A-Xenopus_laevis     LTININILLPPQFPQDKPHISVFPPIHHHLVDKQGMNVIFPPVNNFTMHSDLGKIVQTLL  115
VPS37A-Homo_sapiens       LTININILLPPQFPQEKPVISVYPPIRHHLMDKQGVYVTSPLVNNFTMHSDLGKIIQSLL  120
VPS37A-Mus_musculus       LTININILLPPQFPQEKPVISVYPPIRHHLMDSQGLYVTSPLVSNFTMHSDLGKIIQSLL  120
                          ***************:** ***:***:***:*.**: *  * *.***********:*:**

VPS37A-Xenopus_laevis     EEFWKNPPTLAPSSGSFPYLFSNPAGPQ  143
VPS37A-Homo_sapiens       DEFWKNPPVLAPTSTAFPYLYSNPSGMS  148
VPS37A-Mus_musculus       DEFWKNPPVLAPTSTTFPYLYSNPGGMP  148
                          :*******.***:* :****:***.*
```

**Extended Data Fig. 6 | Sequence alignment of VPS37A N-terminal from different species.** Two highly conserved regions with bulky hydrophobic amino acids which precede and follow the UEVL domain are highlighted in yellow.

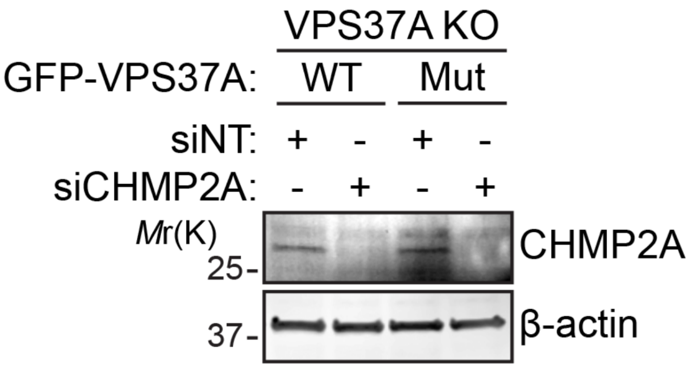


**Extended Data Fig. 7 | Confirmation of siRNA-mediated CHMP2A depletion.** Immunoblot analysis of the indicated siRNA-transfected VPS37A KO U-2 OS cells that were stably transduced with wild-type GFP-VPS37A (WT) or its mutant (Mut, without the first N-terminal 20 amino acid residues and with $F^{137}$PYL to $A^{137}$AAA mutation of VPS37A).

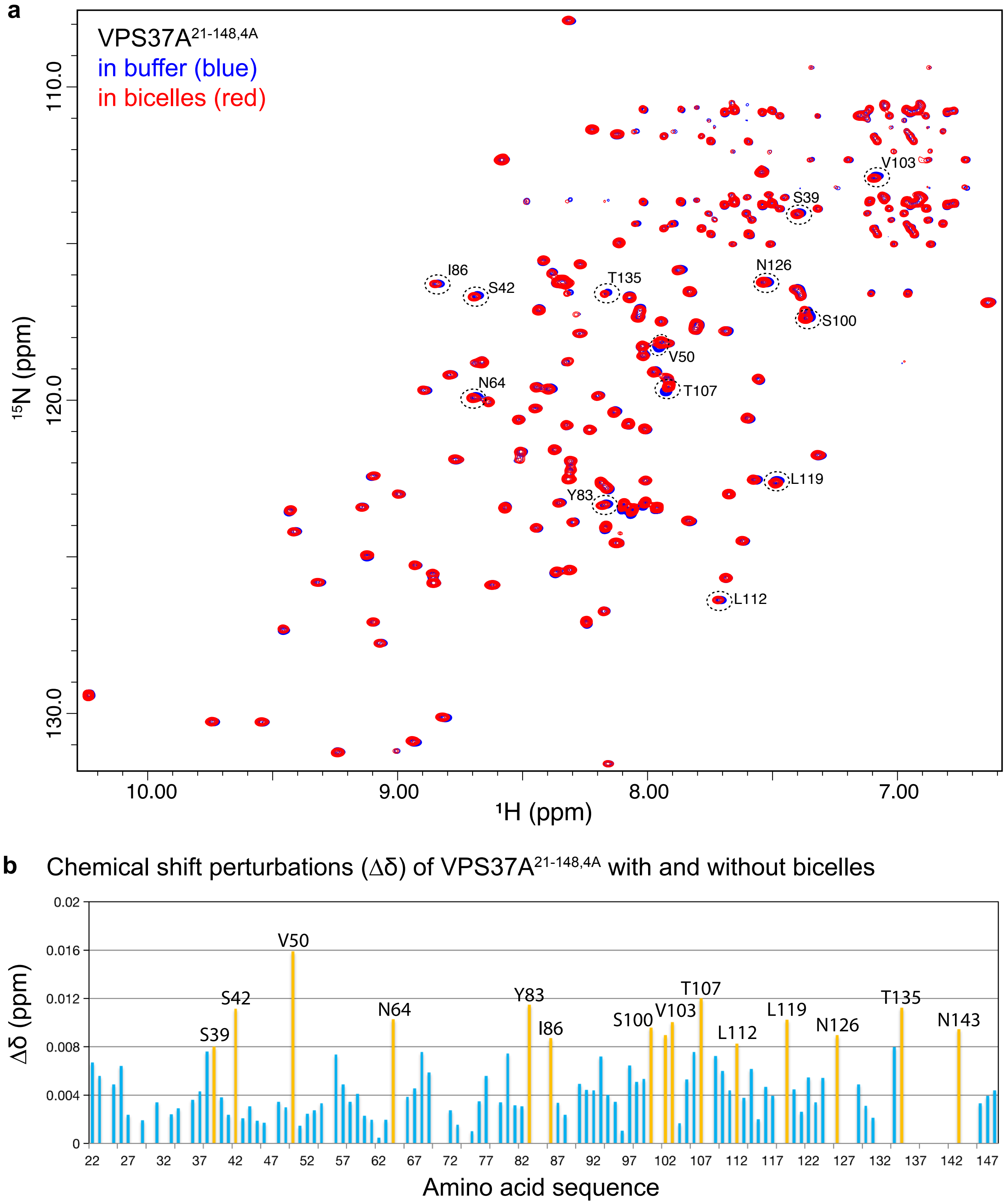


**Extended Data Fig. 8 | VPS37A$^{21\text{-}148,\ 4A}$ membrane interaction.**

**a,** Overlay of $^{15}$N-labeled VPS37A$^{21\text{-}148,4A}$ TROSY spectra acquired on a Bruker 600 MHz spectrometer at 15 °C, pH 7.0 without (blue) and with (red) of bicelles (DMPC:DMPG:DHPC = 4:1:10, molar ratio, q=0.5). Multiple resonances show small but distinct perturbations and are labeled with their assginments.

**b,** Plot of VPS37A$^{21\text{-}148,4A}$ $^{15}$N and $^{1}$H chemical shift perturbations (CSPs, Δδ) upon interaction with bicelles. Unperturbed residues (Δδ < 0.008 ppm) are colored cyan and small perturbed residues (Δδ ≥0.008 ppm) colored orange. The CSPs were calculated as

$$\Delta\delta = \sqrt{0.5 * \{(\frac{\delta_N}{5})^2 + \delta_H^2\}}$$

$\delta_H$ and $\delta_N$ represent the changes in $^{1}$H and $^{15}$N chemical shifts, respectively.

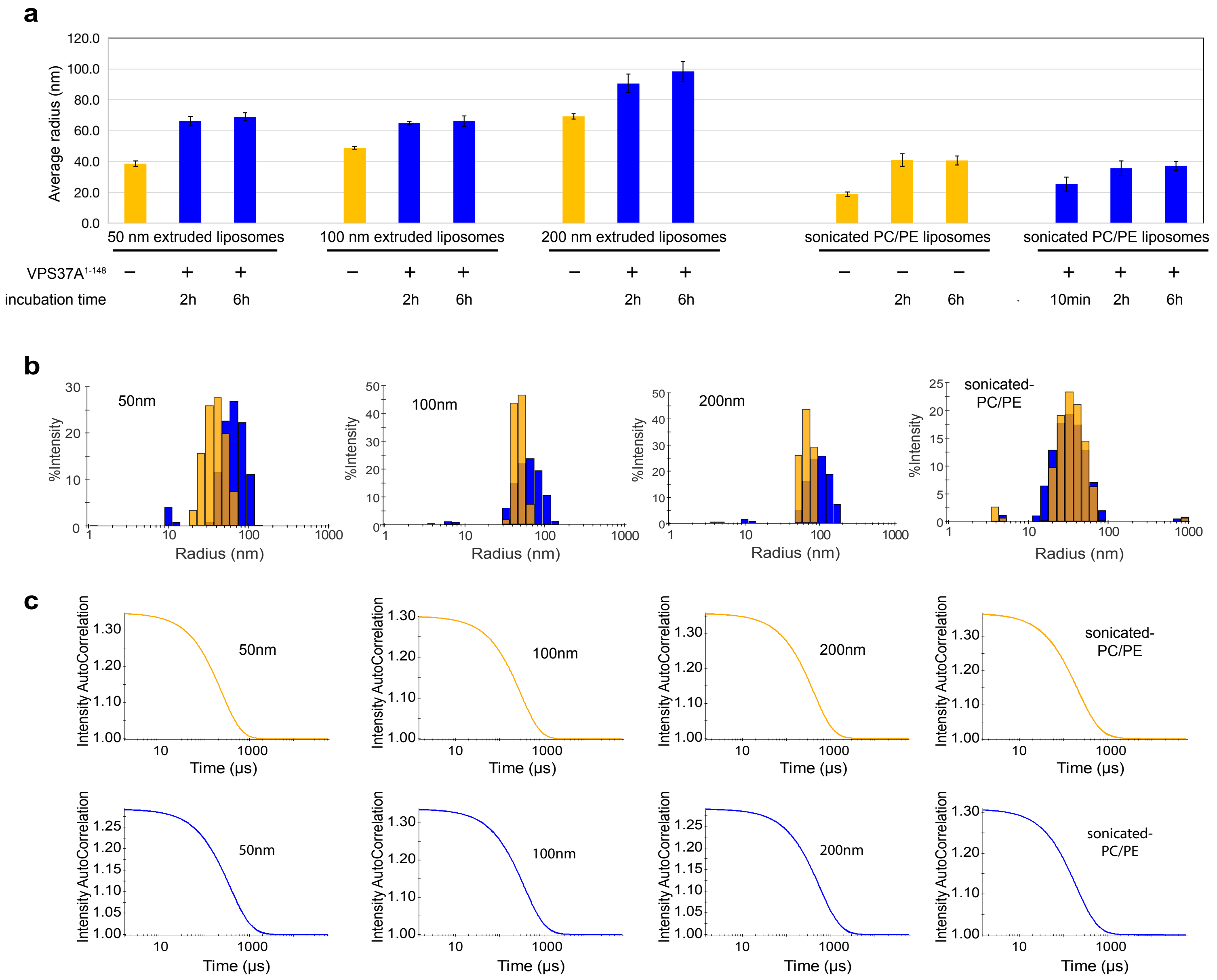


**Extended Data Fig. 9 | Liposome remodeling by VPS37A$^{1-148}$ depends on vesicle size and requires the presence of negatively-charged lipids.**

Extruded liposomes (POPC:DOPG:DOPE=3:2:5) were prepared through membrane with pore sizes of 50nm, 100nm and 200nm. Sonicated liposomes were prepared without PG (POPC:DOPE=45:55, sonicated-PC/PE).

Orange: liposomes only. Blue: liposomes + protein (400:1 molar ratio).

**a,** Average radii of liposomes with and without VPS37A$^{1-148}$ characterized by DLS at indicated incubation times in 50 mM HEPES, pH 7.5, 150 mM NaCl.

**b,** DLS profiles of liposomes with and without VPS37A$^{1-148}$.

**c,** Auto-scaled autocorrelation functions for DLS measurements shown in b.

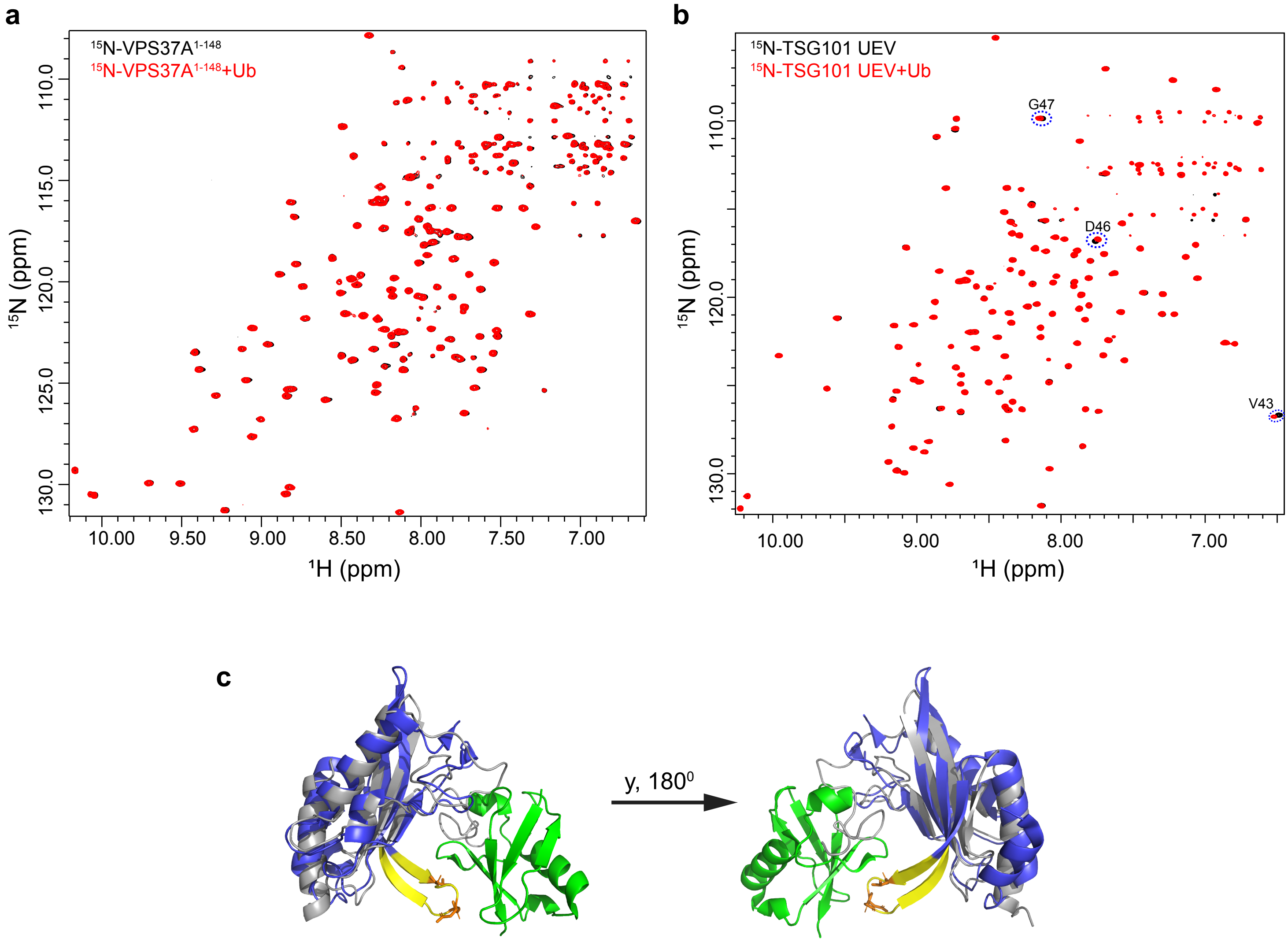


**Extended Data Fig. 10 | VPS37A$^{1\text{-}148}$ domain does not interact with ubiquitin.**
**a,** Overlay of 2D $^{1}$H-$^{15}$N correlation spectra of $^{15}$N-labeled VPS37A$^{1\text{-}148}$ (50 μM) without (black) and with (red) ubiquitin (Ub, 100 μM) in a buffer of 50 mM HEPES, 150 mM NaCl, pH 6.8, and 2 mM TCEP.
**b,** Overlay of 2D $^{1}$H-$^{15}$N correlation spectra of $^{15}$N-labeled TSG101 UEV (50 μM) without (black) and with (red) Ub (100 μM) in a buffer of 50 mM HEPES, 150 mM NaCl, pH 6.8, and 2 mM TCEP. The perturbed resonances are labeled with their assignments.
**c,** Structural alignment of VPS37A$^{21\text{-}148}$ (gray) with TSG101 UEV (blue). The β-tongue of TSG101 UEV is shown in yellow and residues perturbed by the binding of Ub (green) are shown in orange.
